# Generative Access to Make-on-Demand Chemical Space Enables Ultra-Large Virtual Screening

**DOI:** 10.64898/2026.09.17.752345

**Authors:** Kaiyue Zhang, Ying Sun, Xinyue Li, Yuxuan Wang, Chen Peng, Xin Jin, Qi Hu, Jing Huang

## Abstract

Ultra-large-scale virtual screening (ULVS) increasingly relies on make-on-demand chemical spaces, but efficient access to these spaces remains challenging because their scale far exceeds what can be exhaustively enumerated and docked. Here, we present REAL-SWIT, a generative ULVS framework that learns Enamine REAL Space as a synthesizable molecular distribution and couples this generator with a target-specific scoring model to enable docking-guided exploration of REAL Space at the level of complete molecule. The generative model learned transferable features of the REAL Space distribution: approximately 96% of generated molecules were found in REAL Space, and a subset of generated molecules absent from the training release appeared in later expanded releases. In computational benchmarks, REAL-SWIT identified more molecules with favorable docking scores than representative fragment-based search strategies, including cooperative building-block combinations that fragment-level prioritization tended to miss. Experimental validation for ROCK1 yielded six biochemical inhibitors among 23 synthesized compounds, including RX-3 with an IC_50_ of 0.17 µM. These results establish generative access to make-on-demand chemical space as a practical strategy for ULVS.

## 1. INTRODUCTION

Virtual screening (VS) is a central component of early-stage drug discovery, enabling hit identification from ever-expanding chemical libraries. Recent studies have shown that VS exhibits a non-linear dependence on library size, with hit rates increasing abruptly once certain size thresholds are crossed^1,2^. This trend has been confirmed in multiple subsequent large-scale VS campaigns^3–8^. As virtual compound repositories continue to expand^9–11^, VS is increasingly confronted with a fundamental scaling challenge. When library sizes reach the tens-of-billions scale, conventional brute-force pipelines become impractical, owing not only to the computational cost of docking or scoring, but also to the overhead associated with exhaustive enumeration, molecular preparation, and conformer generation. These limitations motivate the development of new algorithmic paradigms for ultra-large virtual screening (ULVS).

The nature of chemical libraries has also undergone a fundamental shift as their size has expanded to the ultra-large regime. Rather than fixed collections of in-stock compounds, modern ultra-large libraries are increasingly defined as make-on-demand, combinatorial chemical spaces, with recent ZINC releases largely drawing from catalogs such as the Enamine REAL library^11^. Advances in parallel computing have made docking at this scale tractable, and screen efficiency can be further improved by machine learning (ML) algorithms^12,13^. In particular, active learning has emerged as an effective strategy, in which docking is performed iteratively on a small subset of compounds to train surrogate models that guide subsequent selection from the library^14–17^. However, these approaches continue to rely on explicit molecular enumeration and do not leverage the combinatorial nature of make-on-demand chemical spaces.

A direct way to move beyond explicit enumeration is to operate in fragment or synthon spaces defined by reagents and their associated reaction protocols^18–22^. V-SYNTHES docks synthons with proper caps, selects fragments that bind favorably, and iteratively expands them geometrically into complete compounds using predefined reaction templates^23,24^. The chemical space docking algorithm docks individual building blocks first and enumerates full compounds only from fragments with favorable interactions^25^. In addition, heuristic search strategies such as Monte Carlo tree search (MCTS) and genetic algorithms have been applied to navigate synthon spaces more adaptively^26–30^. A common feature of these approaches is that fragments must be capped during docking to avoid artifacts from reactive groups; however, the choice of caps is nontrivial and can substantially bias fragment–protein interactions^31^. Furthermore, when fragments are assembled, docking poses and binding energetics can differ from those inferred at the fragment stage. This discrepancy reflects a well-recognized limitation of fragment-based drug discovery, in which favorable fragment-level binding does not necessarily translate into optimal binding of the complete molecule^32^.

In parallel, structure-based molecular generation methods have been developed to directly generate ligands conditioned on target protein pockets. Representative approaches such as Pocket2Mol and FLOWR incorporate structural information through learned geometric priors or structure-aware generation policies^33,34^. The SWIT framework adopts a distinct approach by embedding three-dimensional structural information into molecular generation through docking^35^. By identifying a target-specific favorable subspace of chemical space and learning a docking-based surrogate model within this restricted manifold, SWIT enables targeted generation of molecules with highly favorable docking scores^35^. However, these molecules are often synthetically challenging or inaccessible, severely limiting the practical utility of structure-based molecular generation.

A logical extension is to operate structure-based molecular generation not in the full, unconstrained chemical space but within a conditioned, synthesizable subspace. While synthesizability is difficult to define in general, make-on-demand libraries operationalize it through validated reaction schemas and compatible building-block combinations, which define a vast yet structured chemical manifold that can, in principle, be learned by molecular generative models^36^. In this view, candidate make-on-demand molecules with favorable docking scores can be generated directly, bridging ultra-large virtual screening and structure-based molecular generation. Essentially, generation within a learned synthesizable manifold can be equivalent to performing ULVS over the corresponding combinatory library, but without explicit molecular enumeration.

In this work, we introduce REAL-SWIT, a generative VS paradigm realized within an ultra-large make-on-demand chemical space (Figure 1). We first show that a generative model can be trained to learn the underlying distribution of REAL space with high fidelity, achieving approximately 96% consistency with the combinatorial library. By integrating with docking-based guidance, REAL-SWIT directly identifies REAL Space compounds with favorable interactions for the target pocket. We validate the approach computationally across multiple targets and benchmark it against representative fragment-based ULVS methods. For ROCK1, experimental validation through biochemical and cellular assays shows that REAL-SWIT identifies multiple highly potent hits with novel molecular scaffolds.

**Figure 1.**
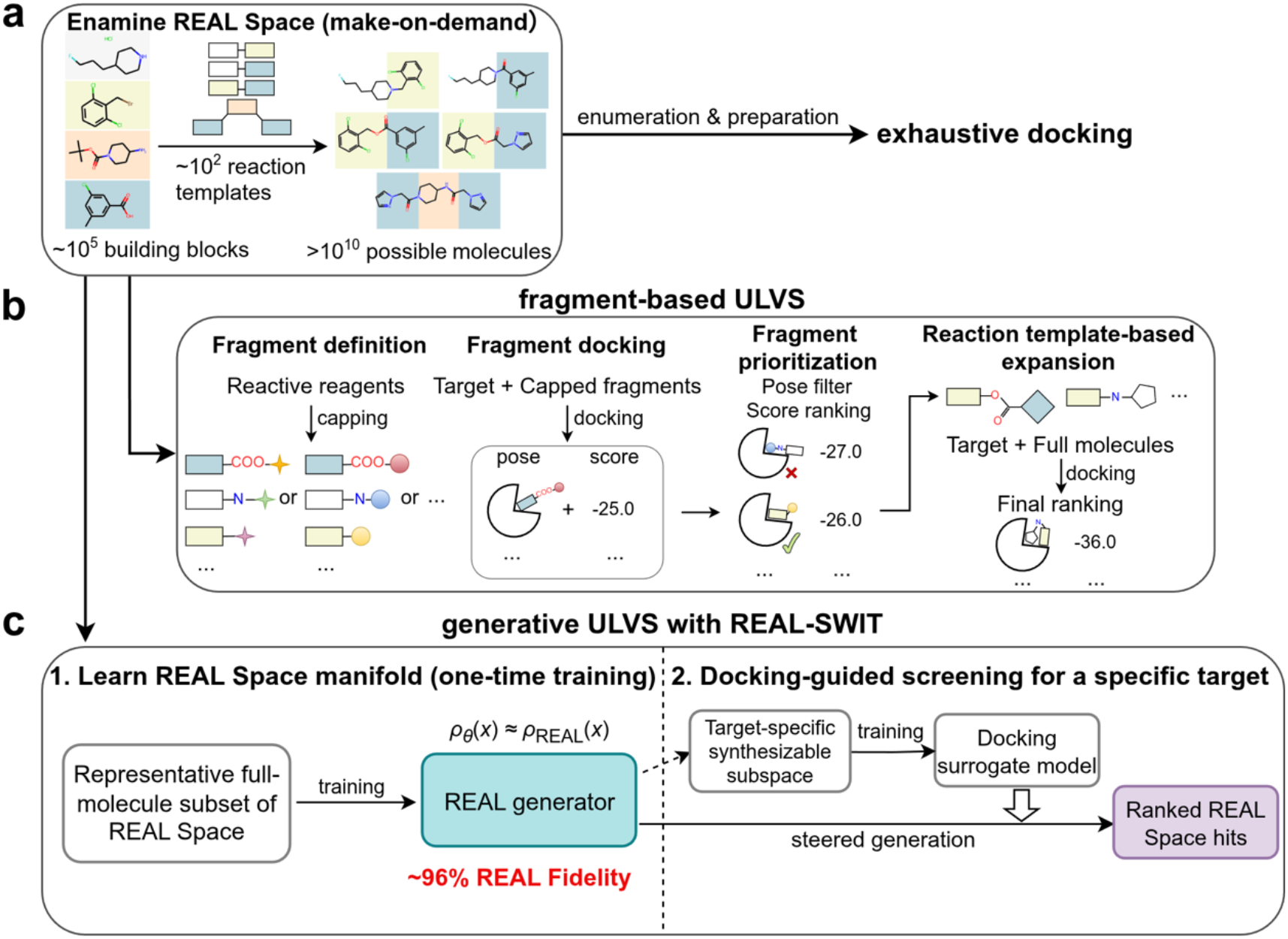
Representative strategies for virtual screening in ultra-large combinatorial chemical spaces. (a) Conceptual illustration of a make-on-demand combinatorial chemical space exemplified by Enamine REAL Space. Vast numbers of synthetically accessible molecules can, in principle, be assembled from available building blocks through validated reaction templates. At this scale, exhaustive enumeration followed by docking requires substantial computational resources. (b) Fragment-based ULVS. Reactive handles of building blocks are capped to generate chemically appropriate fragments for docking. Docked fragments are then prioritized according to pose quality, docking score, and compatibility with template-based expansion. Selected fragments are expanded into full molecules according to predefined reaction templates, and the resulting molecules are subsequently docked to obtain the final target-specific ranking. (c) Generative ULVS with REAL-SWIT. A representative full-molecule subset of REAL Space is curated to train a generative model that learns the underlying manifold of the combinatorial space, enabling direct generation of REAL Space compounds without explicit enumeration. For a given target, a docking surrogate model is trained within a target-specific synthesizable subspace derived from the generator, steering generation toward candidates with favorable predicted docking scores. The output is a ranked list of REAL Space hits for downstream selection for synthesis and experimental testing.

## 2. RESULTS

### 2.1 Learning REAL Space as a generative distribution

REAL-SWIT is designed to perform structure-based VS directly within ultra-large make-on-demand chemical spaces without explicit molecular enumeration. The framework consists of two separable components (Figure 1c): a target-independent generative model that learns the underlying distribution of REAL Space, and a target-specific procedure that trains and uses a docking surrogate model to steer molecular generation toward compounds with favorable interactions for a given binding pocket. A central question underlying this framework is therefore whether the combinatorial structure of REAL Space can be faithfully represented as a learnable generative distribution using only a minute fraction of the library.

Because REAL Space is defined by building blocks and reaction templates rather than by a fully enumerated molecular collection, training such a generative model required constructing representative subsets that could be accessed and processed efficiently. We therefore used a query-based strategy to sample molecules from REAL Space through similarity search (Figure 2a). ChEMBL ligands were used as bioactive-like queries, and for each query we retrieved the top-k most similar REAL molecules, with k = 1, 100, and 1000 yielding three progressively larger training sets of 1.4M, 4M, and 23M molecules after filtering. Generative models trained on these subsets are denoted QB-S, QB-M, and QB-L, respectively (Figure S1). As a size-matched baseline, we trained the same generative architecture on 23M randomly sampled enumerated REAL molecules from ZINC-22, denoted RS.

**Figure 2.**
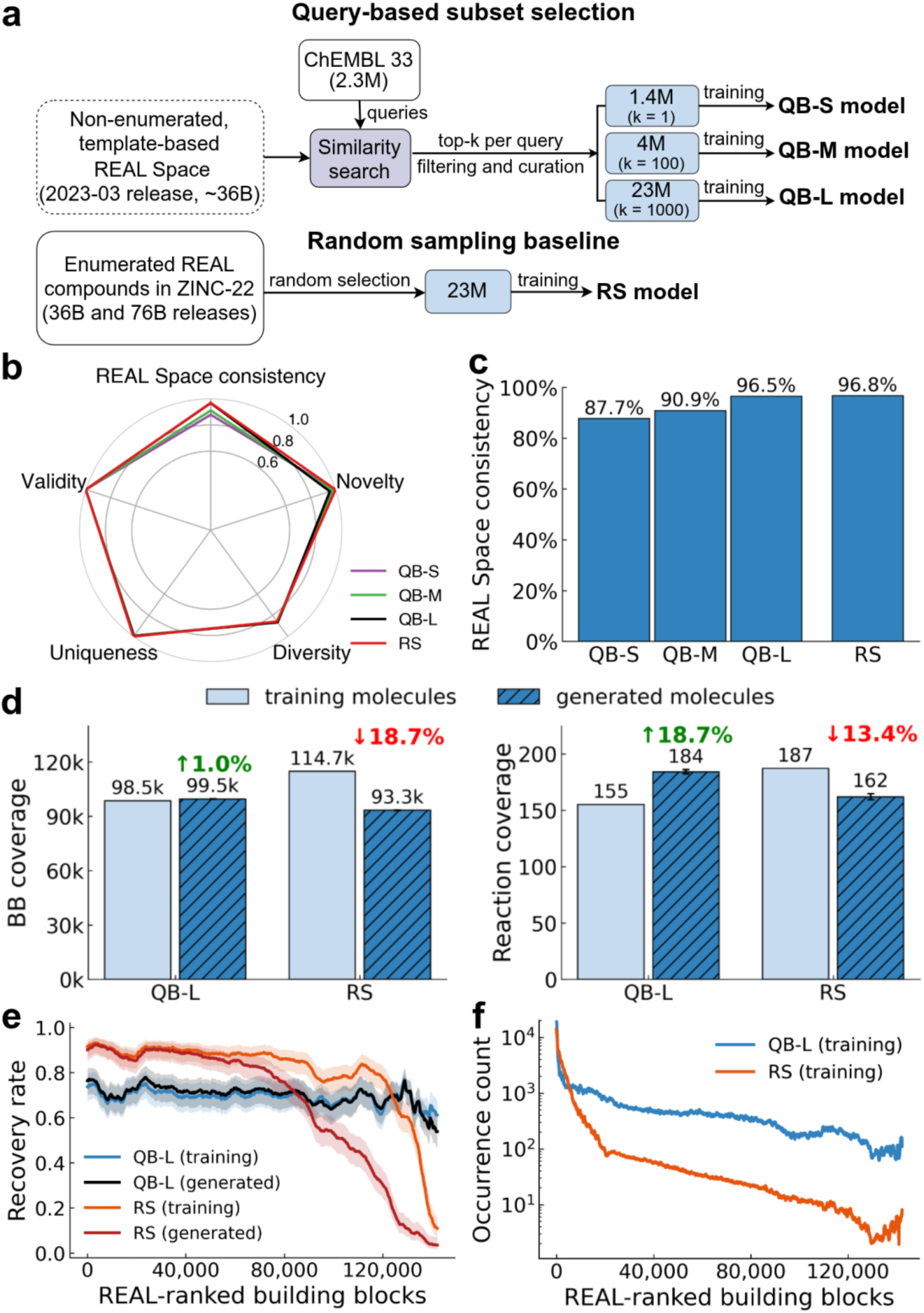
Training generative models from representative subsets of REAL Space. (a) Workflow for constructing representative REAL Space training subsets. Similarity searches using ChEMBL molecules as queries were performed against the non-enumerated, template-based REAL Space library using SpaceLight^20^. (b) Radar plot reporting validity, uniqueness, internal diversity, novelty, and REAL Space consistency for the four models. For each model, values are averaged across five independent sampling runs, with 5 million molecules generated per run. (c) REAL Space consistency of molecules generated by four models, calculated as the fraction of generated molecules found in any of the examined REAL Space releases (36B, 76B, or 83B) and averaged across the five runs. (d) Coverage of REAL Space BBs and reaction types in the training and generated datasets of the QB-L and RS models. For generated datasets, bars and error bars indicate the mean and standard deviations across five independent runs. Percentages reporting the relative change in component coverage in the generated set compared with the corresponding training set are also shown. (e) Recovery rate of REAL Space BBs in the training and generated datasets of the QB-L and RS models as a function of REAL Space occurrence-frequency rank. For each model, one of the five independent generation runs was randomly selected for analysis. Recovery was calculated within BB rank bins of 200 and smoothed using a rolling average over 20 bins. (f) Occurrence counts of REAL Space BBs in the QB-L and RS training datasets plotted against REAL Space frequency rank. Curves were smoothed using a rolling window of 600.

To evaluate model performance, each trained model was sampled in five independent runs, generating 5 million molecules per run for evaluation. Across all models, the resulting molecules showed near-perfect validity and uniqueness, comparable internal diversity, and high novelty (Figure 2b). While these routine generative metrics were uniformly high, we focused on consistency, defined as the fraction of generated molecules present in REAL Space, as the key measure of whether generation remained aligned with the make-on-demand library. We note that the consistency increased progressively with the scale of the query-based training set, reaching 96.5% for QB-L, which was trained on 23M REAL molecules (Figure 2c). The size-matched RS model achieved a similarly high REAL Space consistency of 96.8%. Although 23M molecules are relatively large for molecular generative modeling, approximately 16 times larger than the training set used in REINVENT^37^, it still represents only a minute fraction of REAL Space. Our results show that generative models trained on sufficiently large and diverse subsets can sample predominantly within the combinatorial make-on-demand library. For generated molecules not found in REAL Space, similarity to the nearest REAL Space neighbor remained high, with a median value of approximately 0.75 (Figure S3), suggesting that some of the out-of-library molecules may still lie close to the REAL Space distribution.

REAL Space is not a single static molecular collection but an evolving make-on-demand chemical space, with successive releases expanding as available building blocks and reaction templates are updated. This versioned structure provided an additional test of whether the learned distributions generalize beyond the specific molecular collection used for training. The QB models were trained using molecules retrieved from the 36B REAL Space release (2023-03), whereas consistency was evaluated considering also the expanded 76B (2025-03) and 83B (2025-09) releases. The total in-library consistency values of 87.7%, 90.9%, and 96.5% for QB-S, QB-M, and QB-L, respectively, included contributions from later releases: 4.1%, 3.7%, and 1.8% of generated molecules were absent from the 36B release but present in the 76B release, and 0.4%, 0.4%, and 0.1% were present in the 83B release (Table S1). For the RS model, which was trained on randomly sampled enumerated molecules from a union of the 36B and 76B releases, 0.2% of its in-library consistency was correspondingly attributed to the 83B release.

We further examined the coverage of chemical components and found that QB-L outperformed the size-matched RS model despite their similar consistency. At 5 million generated molecules, QB-L covered 99.5K REAL Space building blocks (BBs) and 184 reaction types, compared with 93.3K BBs and 162 reaction types for RS (Figure 2d). Interestingly, this difference showed opposite trends relative to their respective training sets: the components covered by in-library molecules generated by QB-L exceeded those in its training set (98.5K BBs and 155 reaction types), whereas those from RS remained below its training set (114.7K BBs and 187 reaction types). Thus, QB-L not only recovered but also extrapolated beyond its training-set components. Cumulative recovery curves revealed a similar difference in sampling efficiency (Figure S4). While the numbers of recovered BBs and reaction types approached plateaus by 5 million generations, QB-L had already recovered 93.7K BBs and 178 reaction types at 1 million generations, compared with a slower accumulation of 64.8K and 144 for RS.

To understand the origin of this difference, we compared BB recovery as a function of REAL occurrence frequency. We ranked BBs by their occurrence frequency in REAL Space and compared their recovery in the training and generated molecules of QB-L and RS (Figure 2e). Molecules generated by QB-L closely followed the BB recovery profile of the QB-L training set across the frequency range, including the lower-frequency tail, whereas molecules generated by RS showed a pronounced loss of lower-ranked BBs. Further inspection on the training sets revealed distinct BB occurrence distributions induced by the two subset construction strategies (Figure 2f). Query-based sampling enriched lower-frequency BBs and produced a more balanced training distribution, with many tail BBs still appearing close to 100 times and only 1.6% of BBs represented fewer than 10 times. In contrast, random sampling displayed a highly skewed BB occurrence distribution, with many BBs occurring only rarely; specifically, 24.4% of BBs appeared fewer than 10 times. This result suggests that sufficient representation of lower-frequency BBs, rather than their mere inclusion in the training set, is critical for learning the component-level structure of REAL Space. Taken together, these analyses show that query-based subset construction supports high in-library consistency, extension to later REAL Space releases, and efficient recovery of BB and reaction-type coverage, establishing QB-L as the generative foundation for subsequent REAL-SWIT screening experiments.

### 2.2 REAL-SWIT expands access to high-scoring molecules beyond fragment-based search

We next integrated the QB-L model into the structure-based SWIT framework for generative ULVS. In this target-specific stage, REAL-SWIT trains docking surrogate models to approximate docking scores and steer full-molecule generation toward candidates with more favorable predicted scores (Figure 1c). A key finding from our original SWIT work was that, to maintain surrogate transferability in predicting docking scores during target-directed generation, surrogate training should be performed in a two-stage manner^35^. In REAL-SWIT, we first docked 100,000 randomly generated REAL molecules to form an initial random set (R1), which was used to train a target-specific D-MPNN scoring model. This surrogate model was then used to guide a preliminary generation round, producing 1 million molecules, from which the top 100,000 ranked molecules were selected as the first target-specific set (V1) and subjected to docking calculations. Because R1 samples REAL Space broadly whereas V1 is enriched toward the target, we combined R1 and V1 to define a target-specific synthesizable subspace and trained another surrogate model, with improved correlation between predicted and docking scores for generated molecules against the specific target^35^. This refined model guided a second generation round of 1 million molecules, which were further screened by SpaceLight to verify their presence in REAL Space. The top 100,000 retained molecules according to surrogate-predicted scores were then selected for docking and constituted the final output set (V2) for downstream analysis.

We benchmarked REAL-SWIT on two targets, cannabinoid receptor 2 (CB2) and Rho-associated protein kinase 1 (ROCK1), using ICM-Pro^38^ for all docking calculations. Both the targets and docking settings matched those used in the V-SYNTHES study^23^, enabling direct comparison with this representative fragment-based ULVS method. We further implemented SyntheMol^28^, which utilizes MCTS to explore combinatorial chemical space by treating BBs as child nodes and expanding candidate molecules according to node scores and pre-defined reaction rules. In our implementation, initial node scores were obtained from ICM-Pro docking, and complete molecules generated during rollout were evaluated using the same target-specific docking surrogate model as in REAL-SWIT. One million compounds were generated, and the top-ranked 100,000 were selected for docking. Together, these two fragment-based search methods provided strong baselines for assessing the performance of REAL-SWIT.

Across both targets, REAL-SWIT substantially enriched for REAL Space compounds with more favorable docking scores (Figure 3a, b). The docking-score distribution of molecules generated by REAL-SWIT shifted toward more favorable values compared with V-SYNTHES and SyntheMol, with the difference most pronounced at stringent score thresholds. REAL-SWIT identified 10 compounds with ICM-Pro docking scores below −50 for CB2 and 106 compounds for ROCK1, whereas the fragment-based methods yielded few or no compounds in this range. At a docking-score cutoff of −40, REAL-SWIT identified 2,438 and 12,341 compounds for CB2 and ROCK1, respectively. In contrast, SynthMol identified 304 and 876 compounds, whereas V-SYNTHES reported 29 and 2,521 compounds, respectively. The most favorable docking scores reported in the V-SYNTHES study were −44.0 for CB2 and −49.5 for ROCK1. We note that redocking of the 28 CB2 active molecules reported in the V-SYNTHES study revealed a modest discrepancy between our calculated scores and the published values, with a mean absolute difference of 0.7 (Figure S5), potentially reflecting slight differences in ICM-Pro settings and stochasticity in docking calculations. If we considered −30 as the docking-score cutoff for hits, 62.5% of CB2 compounds and 90.1% of ROCK1 compounds in the REAL-SWIT V2 sets were classified as hits, compared to 2.3% and 1.0% in the randomly generated R1 set. This corresponds to 27- and 87-fold enrichment for CB2 and ROCK1, respectively, broadly consistent with the 39- and 42-fold enrichment reported for V-SYNTHES over standard VS^23^.

**Figure 3.**
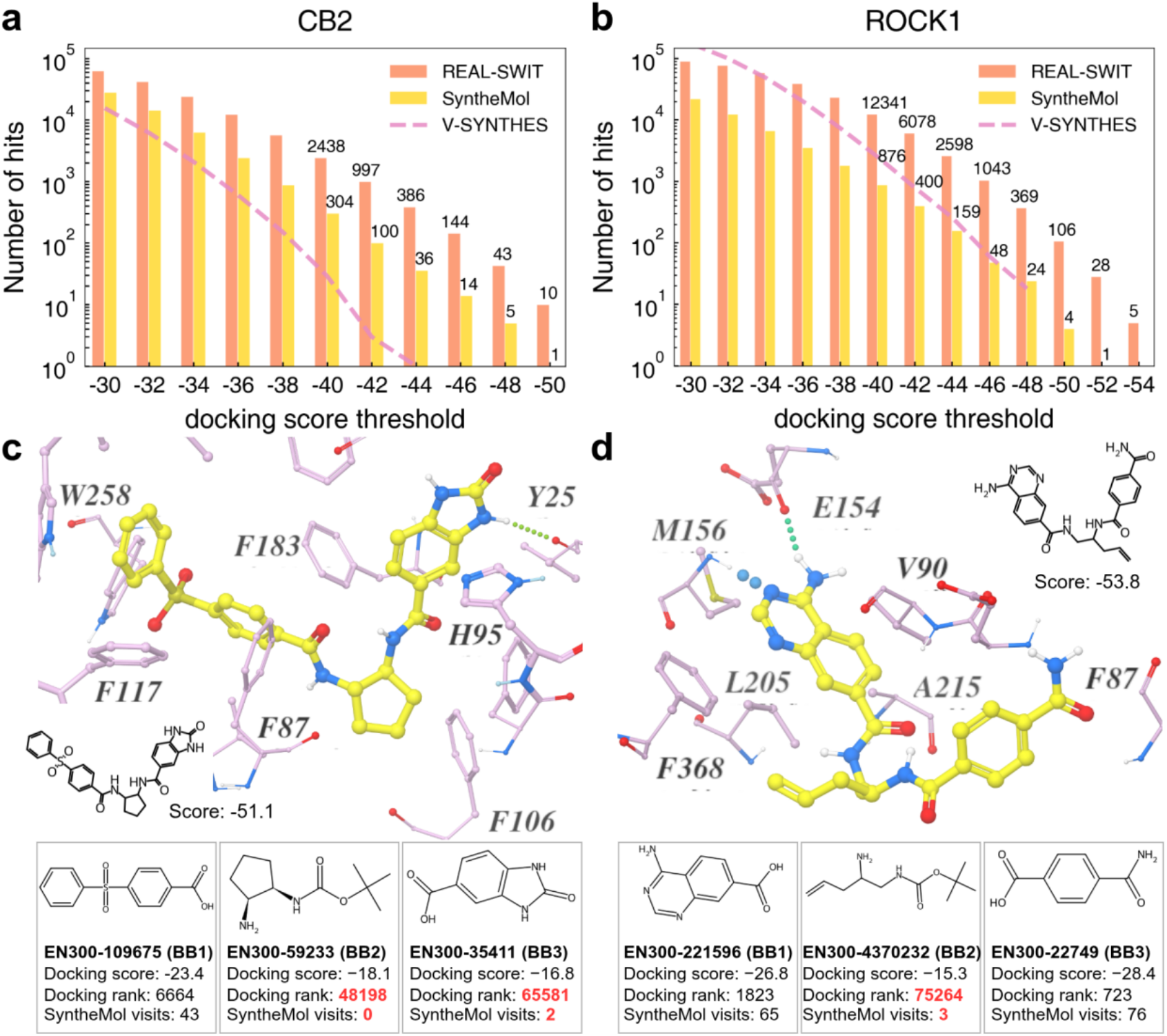
Comparison of screening performance between REAL-SWIT and fragment-based ULVS methods. (a-b) Numbers of hits with docking scores below progressively less stringent thresholds among molecules generated for (a) CB2 and (b) ROCK1. The top 100,000 compounds from REAL-SWIT (orange) and SyntheMol (yellow) were docked under the same conditions and compared with V-SYNTHES results (pink) reported in the original study. For V-SYNTHES, the CB2 results combine outputs from the screened two- and three-component reaction libraries, with 1.5 million compounds docked in total, whereas the ROCK1 results were based on 1.0 million docked compounds from the two-component reaction library only. (c-d) Representative high-scoring molecules identified by REAL-SWIT for (c) CB2 and (d) ROCK1 that were not recovered by SyntheMol, shown with their chemical structure, docking poses and docking scores. Constituent BBs are annotated with their docking scores, fragment ranks, and visit counts during the selection phase of the SyntheMol search.

To understand why fragment-based search recovered fewer high-scoring REAL Space molecules, we examined how the BBs composing high-scoring REAL-SWIT hits were treated during the SyntheMol search process. This analysis showed that, because SyntheMol prioritizes BB nodes using fragment-level docking scores, BBs that score weakly in isolation may be sampled only rarely, even if they become important in the assembled molecule. Two illustrative examples are shown in Figure 3. For CB2, REAL-SWIT identified a three-component compound with a docking score of −51.1 (Figure 3c). While BB1 had a relatively favorable docking score and was visited 43 times by SyntheMol, BB2 and BB3 as fragments had weaker docking scores, lower ranks, and were visited 0 and 2 times during the selection, respectively. In the representative ROCK1 hit (Figure 3d), the two terminal BBs were favorably ranked and frequently visited, whereas the linker BB ranked much lower and was visited only 3 times, despite being part of a REAL molecule that docks very favorably as a whole.

These examples highlight a key limitation of fragment-level prioritization: the contribution of a BB may depend strongly on its context in the complete molecule rather than on its isolated docking pose and score. This effect is particularly clear for linker-like BBs, which may not dock favorably on their own but can orient other components and enable a favorable full-molecule pose. Consistent with this view, we found that a majority of REAL Space molecules generated by SyntheMol for both targets were two-component products, suggesting systematic underexploration of linker-containing combinations. A direct comparison between independently docked fragments and their corresponding full-molecule poses further showed that fragment poses can deviate substantially from their positions in the assembled molecules (Figure S6). Similar fragment-to-molecule pose deviations were also reported in the V-SYNTHES2 study for three-component molecules targeting rhodopsin^24^. Interestingly, in the case of ROCK1, independently docked BBs preferentially occupied the kinase hinge region, potentially biasing fragment-level sampling toward locally favorable hinge-binding poses while reducing exploration of other positions required for productive full-molecule assembly. By generating complete molecules directly, REAL-SWIT can therefore identify cooperative BB combinations that are less readily captured by fragment-based search.

### 2.3 REAL-SWIT performs target-guided broad-to-local search in combinatorial chemical space

Beyond the final output sets, the generation trajectories of REAL-SWIT can be analyzed explicitly in terms of the BBs and reaction types that define REAL Space. We therefore first examined how rapidly target-guided generation explored these chemical components during the second generation round (Figure 4a). For both CB2 and ROCK1, the cumulative numbers of unique BBs and reaction types increased sharply during the early phase and approached a plateau after approximately 2,000 steps. By this stage, REAL-SWIT had sampled more than half of the BBs present in the QB-L training set and nearly all reaction types, while many of the components appearing in the final top-ranked 100,000 molecules had also already been encountered.

**Figure 4.**
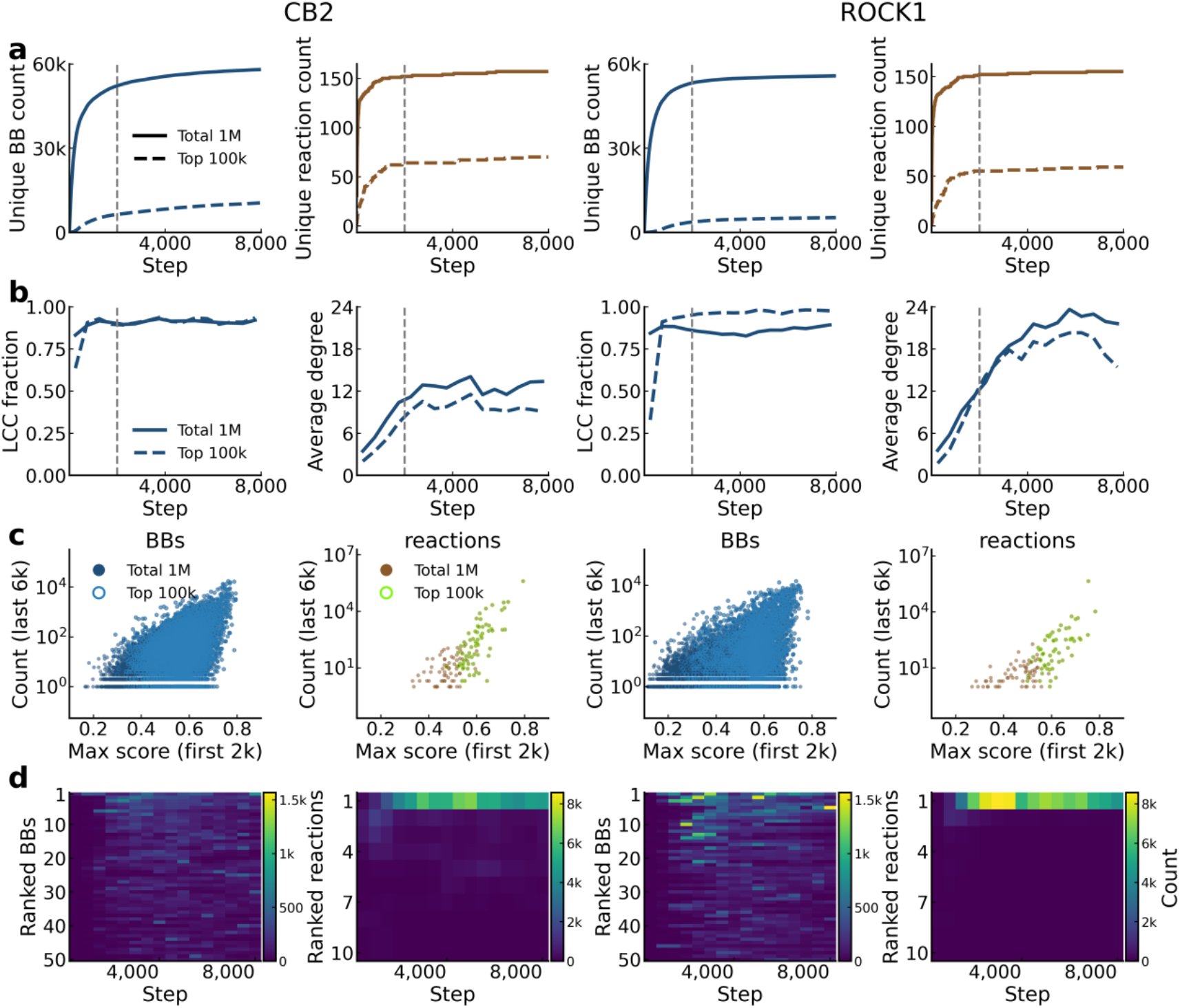
Analysis of the REAL-SWIT generation processes. (a) Cumulative recovery of unique BBs and reaction types during REAL-SWIT generation. Solid lines represent the full generated set of 1 million molecules, whereas dashed lines represent molecules retained in the final top-ranked 100,000 molecules. (b) Evolution of BB co-occurrence network connectivity. Generated molecules were grouped into 500-step bins, and a BB co-occurrence network was constructed for each bin. In each network, BBs were represented as nodes, with an edge added between each pair of BBs co-occurring within the same molecule. The fraction of BBs contained in the largest connected component and the average degree of the network were calculated to characterize the extent and density of BB recombination, respectively. (c) Relationship between early component-level scoring and later component usage. For each BB or reaction type observed during the first 2,000 generation steps, the x-axis shows its maximum normalized D-MPNN score among molecules containing that component within this early-generation window, and the y-axis shows its total occurrence during the last 6,000 generation steps. Components retained in the final top-ranked 100,000 molecules are highlighted with open circles. (d) Temporal usage patterns of the most frequently sampled components. Heatmaps show the occurrence counts of the top 50 BBs and top 10 reaction types, ranked by their overall frequencies in the final top-ranked molecules.

We also examined how these sampled components were combined during generation. To capture the evolving connectivity of the search trajectory, we constructed BB co-occurrence graphs at 500-step intervals, in which BBs were represented as nodes and their co-occurrence within generated molecules as edges (Figure 4b). For both targets, the fraction of BBs in the largest connected component (LCC) increased rapidly and remained high after the early phase, indicating that generated molecules became organized around an emerging manifold of BB combinations. The average degree of BBs also increased over the generation trajectory, showing that sampled BBs were increasingly recombined with one another. Together with the early saturation of component coverage, this increasing connectivity supports a broad-to-local search pattern, in which REAL-SWIT first establishes broad access to the chemical components of the combinatorial space and then performs denser recombination within target-specific subspaces.

We next analyzed the relationship between the early scoring of chemical components and their later sampling during generation. For each BB and reaction type, we calculated an early max score, defined as the maximum surrogate-predicted score among molecules containing that component during the first 2,000 generation steps (Figure 4c). The scores were normalized using sigmoid function (Eq. 2), such that higher normalized values corresponded to more favorable predicted docking scores. We then compared this component-level score with the occurrence count of the same BB or reaction type during the last 6,000 generation steps. For both targets, components with higher early max scores tended to be sampled more frequently later in the trajectory, consistent with progressive amplification of target-favored regions in the combinatorial space. However, this prioritization was not exclusive: many components with moderate early scores continued to be sampled, and components contributing to the final top-ranked 100,000 molecules spanned a broad range of early scores. Thus, REAL-SWIT realizes a soft target-guided prioritization scheme, biasing generation toward promising components while preserving sufficient diversity for later recombination.

Finally, we examined the temporal usage patterns of the most frequently sampled BBs and reaction types throughout generation (Figure 4d). The heatmaps showed that REAL-SWIT did not immediately collapse onto a fixed set of components during the search. Instead, frequently sampled BBs continued to turn over across the trajectory, while reaction-type usage became more focused, with a smaller number of reaction types repeatedly used at later stages. Together with the rapid saturation of component coverage, the formation of an emerging manifold of BB combinations, and the component-level relationship between early scores and later usage, these results support a central picture of the REAL-SWIT search process: the model first establishes broad access to chemical components available in the combinatorial space and then progressively performs denser, target-guided recombination within target-specific subspaces.

### 2.4 REAL-SWIT samples neighborhoods surrounding known actives in chemical space

In virtual screening, the primary objective is to identify active-like compounds that can serve as starting points for subsequent optimization. We therefore examined the relationship between REAL-SWIT-generated molecules and known actives for CB2 and ROCK1. For each generated compound, we calculated the maximum Tanimoto coefficient (TC) similarity to known actives for the corresponding target that were present in REAL Space. We collected 6,320 ChEMBL and 73 V-SYNTHES actives for CB2, and 1,607 ChEMBL and 6 V-SYNTHES actives for ROCK1. Among these known actives, 216 CB2 compounds and 136 ROCK1 compounds were present in REAL Space. We docked these in-library actives using the same ICM-Pro protocol and compared their score distributions with those of REAL-SWIT generated molecules. The in-library actives generally showed less favorable ICM-Pro scores than the generated compounds (Fig. 5e–f, upper panels). We do note that this distributional difference does not imply that most generated compounds would be active; rather, it highlights the empirical nature of docking scores and their limited ability to pinpoint experimentally active compounds at the individual-molecule level. In general, docking scoring functions are useful for enriching active compounds, but their quantitative inaccuracies remain a limiting factor in docking-based VS workflows. Our previous work further showed that docking-guided generative models can exploit the artifacts in docking scores^35^.

**Figure 5.**
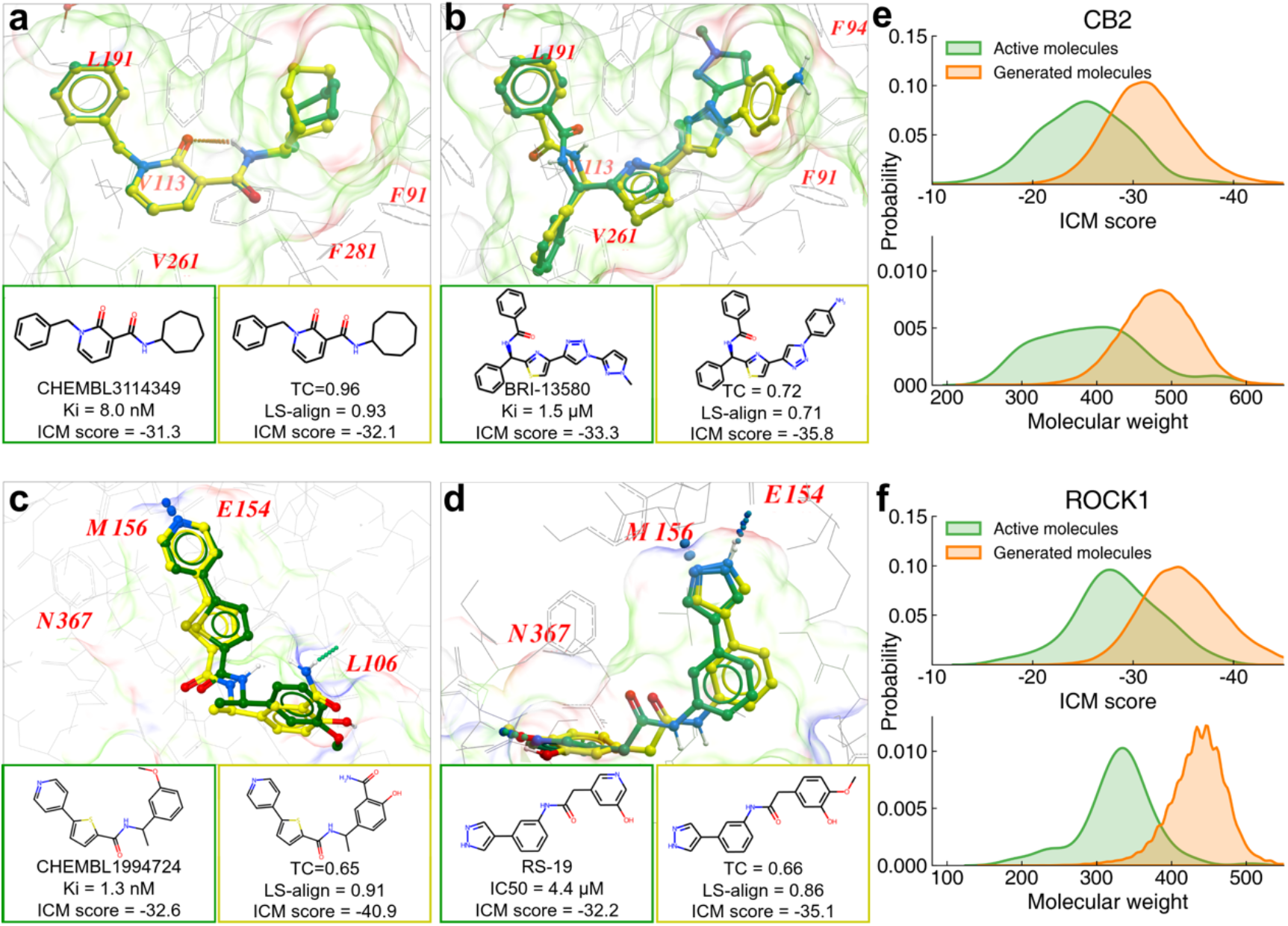
Comparisons of known actives and structurally related REAL-SWIT-generated molecules. (a-d) Structural comparisons of generated compounds (yellow) with known actives (green) for CB2 and ROCK1. Docking poses were obtained using ICM-Pro, 2D similarity was measured by the Tanimoto coefficient (TC), and 3D alignment was evaluated using LS-align^42^. (e-f) Distributions of ICM-Pro docking scores (top) and molecular weights (bottom) for in-library actives and top-ranked 100,000 generated molecules.

Comparison of molecular weights showed that the generated molecules were substantially larger (Fig. 5e–f, lower panels), likely reflecting the enrichment of three-component products that can form more extensive interactions with the binding pocket. In contrast, in-library actives were dominated by two-component REAL Space molecules, accounting for all ROCK1 actives and 90% of CB2 actives. This observation is consistent with the known size dependence of docking scores^39^ and illustrates how docking-based ULVS can amplify empirical features of scoring functions. Accordingly, ligand efficiency and ligand-lipophilicity efficiency have been proposed as alternative optimization metrics^40,41^, which could be straightforwardly incorporated into the current REAL-SWIT workflow as driving targets.

Despite the difference in overall score distributions, REAL-SWIT generated molecules that closely resembled known actives with relatively favorable docking scores. Four representative examples are illustrated in Figure 5, including one ChEMBL-related and one V-SYNTHES-related molecule for each target. For CB2, one generated molecule closely matched the ChEMBL active ChEMBL3114349 (8 nM), with the terminal cycloheptyl group replaced by a cyclooctyl group, while maintaining a similar binding pose and improving the ICM-Pro score from −31.3 to −32.1. Another generated molecule differed with the V-SYNTHES active BRI-13580 by a terminal N-methylimidazolyl group being replaced by para-aminophenyl. This substitution did sample a different interaction pattern and improve the ICM-Pro score. For ROCK1, the generated molecules likewise showed only minor structural differences from ChEMBL1994724 and the V-SYNTHES active RS-19, while retaining similar binding modes and achieving better docking scores. More broadly, REAL-SWIT recovered constituent BBs from a high proportion of curated in-library actives with favorable ICM-Pro scores (≤ −30), covering 87.9% of CB2 actives and 78.7% of ROCK1 actives (Table S3). These result show that REAL-SWIT can effectively sample neighborhoods surrounding known active compounds in the chemical space.

### 2.5 Experimental validation identifies novel ROCK1 inhibitors with biochemical potency and cellular activity

The analyses above show that REAL-SWIT can generate docking-favorable molecules within REAL Space and sample neighborhoods surrounding known actives, but the ultimate goal of ULVS is to identify compounds with experimentally measurable activity. We therefore selected ROCK1 for evaluation, as the same target had previously been used to evaluate V-SYNTHES and Chemical Space Docking, providing relevant reference compounds for comparison^23,25^. The top-ranked 100,000 REAL-SWIT-generated ROCK1 molecules were filtered for drug-like properties and then clustered based on protein-ligand interaction fingerprints^43^. The compound with the best ICM-Pro score from each cluster was subjected to visual inspection, with particular emphasis on forming at least one hydrogen bond with the hinge residue Met156. This prioritization strategy yielded 28 candidates ordered for synthesis from Enamine, of which 23 were successfully delivered and advanced to ROCK1 enzymatic inhibition assays.

ROCK1 inhibitory activity was assessed using a luminescence-based ADP-Glo™ that quantifies ADP formation during catalysis. To benchmark REAL-SWIT against previous ULVS studies on the same target and make-on-demand space, we tested two reported ROCK1 inhibitors, RS-15 from V-SYNTHES and CS-1 from Chemical Space Docking^23,25^. The 23 synthesized REAL-SWIT compounds, together with the two reference compounds, were initially screened at 10 µM (Figure 6a). Six REAL-SWIT compounds reduced ROCK1 activity by more than 30% at this concentration, corresponding to a primary hit rate of 26.1%. Among them, RX-3 was the strongest hit in the single-point assay, reducing residual ROCK1 activity to approximately 5% of the DMSO control and showing stronger inhibition than RS-15, and CS-1 under the same assay conditions.

**Figure 6.**
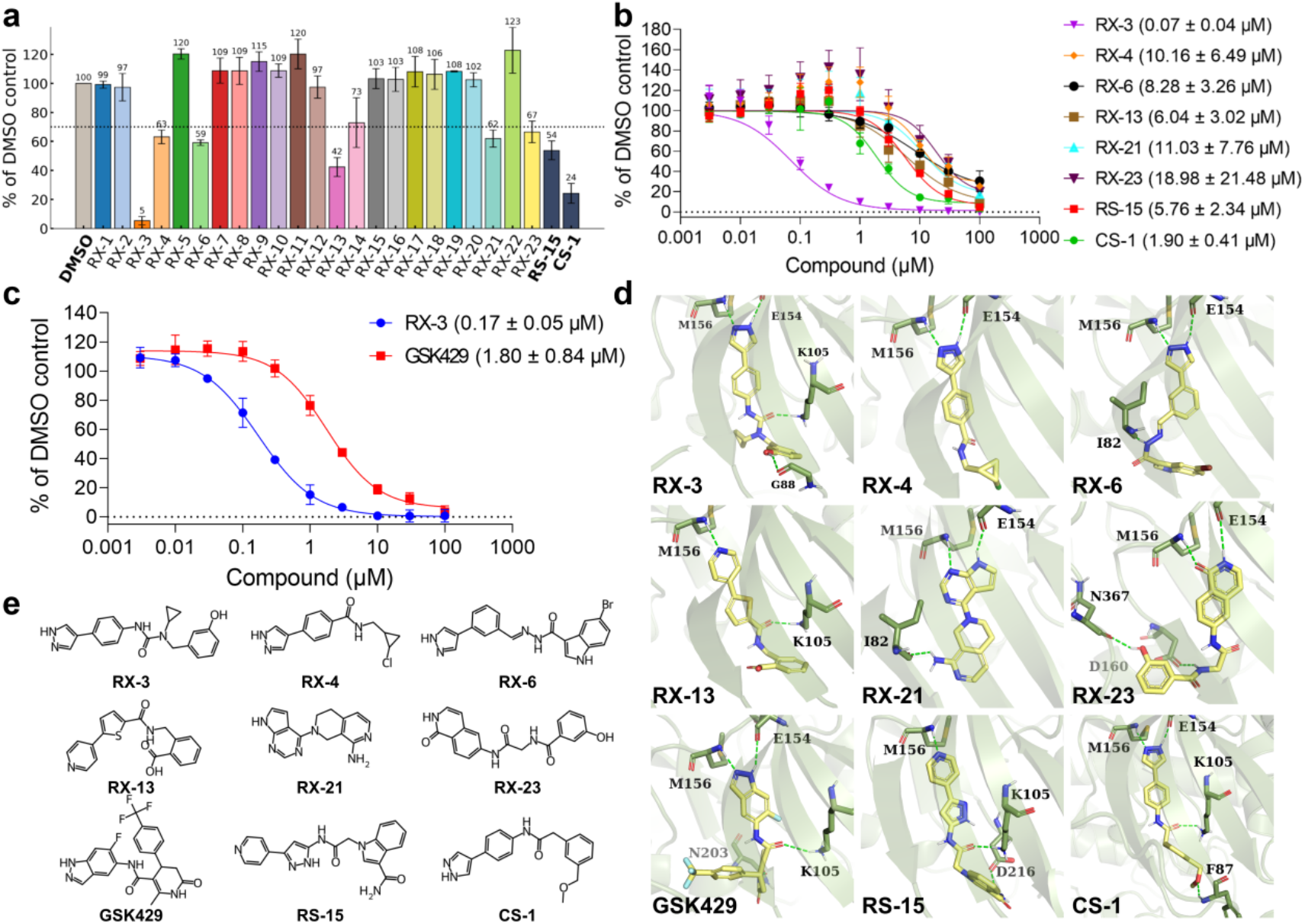
Experimental validation and binding-mode analysis of ROCK1-targeted hits generated by REAL-SWIT. (a) Residual ROCK1 enzymatic activity measured for 23 synthesized compounds and two reference compounds, RS-15 and CS-1. Compounds were tested individually at 10 µM, and activities were normalized to the DMSO control. The dashed line indicates 70% residual activity, corresponding to 30% inhibition. (b) Dose-response curves for six active generated compounds, RS-15, and CS-1. (c) Dose-response curves for RX-3 and GSK429286A (abbreviated as GSK429). (d) Binding poses of nine compounds within the ROCK1 ATP-binding pocket. The binding poses of the generated compounds, GSK429, and RS-15 were predicted using ICM-Pro, whereas that of CS-1 was obtained from its crystal structure (PDB ID: 7S25). Key hinge-binding interactions and additional hydrogen bonds are highlighted. (e) Chemical structures of the nine compounds. Data points are presented as mean ± s.d. from three independent experiments. IC50 values are reported with standard errors from nonlinear regression fitting in GraphPad Prism.

Dose–response assays were then performed for the six REAL-SWIT hits and the two reference inhibitors (Figure 6b). All six REAL-SWIT compounds showed concentration-dependent inhibition of ROCK1, confirming that the primary-screen hits reflected reproducible biochemical activity. RX-3 was the most potent compound in this assay, with an IC_5 0_ of 0.07 ± 0.04 µM, substantially lower than those of RS-15 and CS-1 under the same assay conditions. We note that RX-3 and CS-1 share one common BB but, as two-component REAL Space compounds, differ in the other BB and reaction type. Their binding poses are also similar, with the shared BB forming the same hinge-binding pattern and amide-carbonyl interaction with Lys105, while the nonshared moieties each provide an additional peripheral hydrogen bond, donated to Gly88 in RX-3 and accepted from Phe87 in CS-1. The remaining REAL-SWIT hits showed low-micromolar activity. Among them, RX-13 showed potency (6.04 µM) comparable to RS-15, further supporting the ability of REAL-SWIT to identify experimentally active ROCK1 inhibitors from the generated set.

We further compared RX-3 with GSK429286A, a commercially available and well-established potent ROCK1 inhibitor^44^. In this independent dose-response experiment, RX-3 showed an IC_50_ of 0.17 ± 0.05 µM, approximately tenfold than that of GSK429286A (Figure 6c). Together with the comparison against RS-15 and CS-1, this independent benchmark confirmed RX-3 as a potent ROCK1 inhibitor. We note that apparent IC_50_ values can depend on assay condition, particularly ATP concentration for ATP-competitive kinase inhibitors. The assays here were performed at 10 µM ATP, compared with 1 µM ATP in the previous studies^44^, providing a more stringent biochemical setting and potentially contributing to differences in the absolute IC_50_ values.

We next assessed the novelty and diversity of the six experimentally validated REAL-SWIT hits. To quantify their structural similarity to previously reported ROCK1 inhibitors, each hit was compared against ROCK1 actives from ChEMBL, V-SYNTHES, and Chemical Space Docking using TC similarity (Table S4). All six hits showed maximum similarity values below 0.65, indicating that they are distinct from known ROCK1 inhibitor chemotypes. RX-21 was the most structurally distinct compound, with a maximum similarity of only 0.43 to any reported ROCK1 inhibitor. Consistent with the interaction-fingerprint clustering strategy used during compound prioritization, the six hits also displayed diverse chemical structures and adopted distinct binding poses within the ROCK1 ATP-binding pocket (Figure 6d). These results show that the experimentally confirmed REAL-SWIT hits are not redundant analogs of known inhibitors, but represent multiple novel starting points for subsequent lead optimization.

While ROCK1 is a validated target in cancer-related cellular processes, ROCK1 inhibitors also have broad practical utility as cell-culture reagents, particularly in stem cell recovery and organoid culture^45–47^. We therefore asked whether representative REAL-SWIT-derived ROCK1 inhibitors showed measurable activity in cell-based assays beyond biochemical inhibition. All six REAL-SWIT hits and the reference inhibitors were evaluated in soft-agar colony formation assays using A549 and H1299 cells, with Y-27632 included as a standard ROCK inhibitor widely used in cell-culture applications^45^. Among all tested compounds, RX-21 showed the strongest active generated compound, with IC_50_ values of 5.62 in A549 cells and 9.73 µM in H1299 cells, slightly outperforming Y-27632 under the same assay conditions (Figure 7). In contrast, the most potent biochemical inhibitor RX-3 did not show corresponding cellular activity, highlighting that biochemical potency and cellular activity are not fully coupled (Figure S7). We further measured 2D adherent cell proliferation by real-time live-cell imaging to assess whether the suppression of colony formation was specific or simply reflected general growth inhibition. Confluency profiles showed that, for all three compounds shown in Figure 7 (RX-21, CS-1, and Y-27632), effects on general proliferative growth occurred at substantially higher concentrations than those that reduced colony formation, suggesting that these compounds suppress anchorage-independent growth without broadly inhibiting cell proliferation at the same concentration range. Together, these results show that REAL-SWIT can identify novel ROCK1 inhibitors with measurable activity in functional cellular assays.

**Figure 7.**
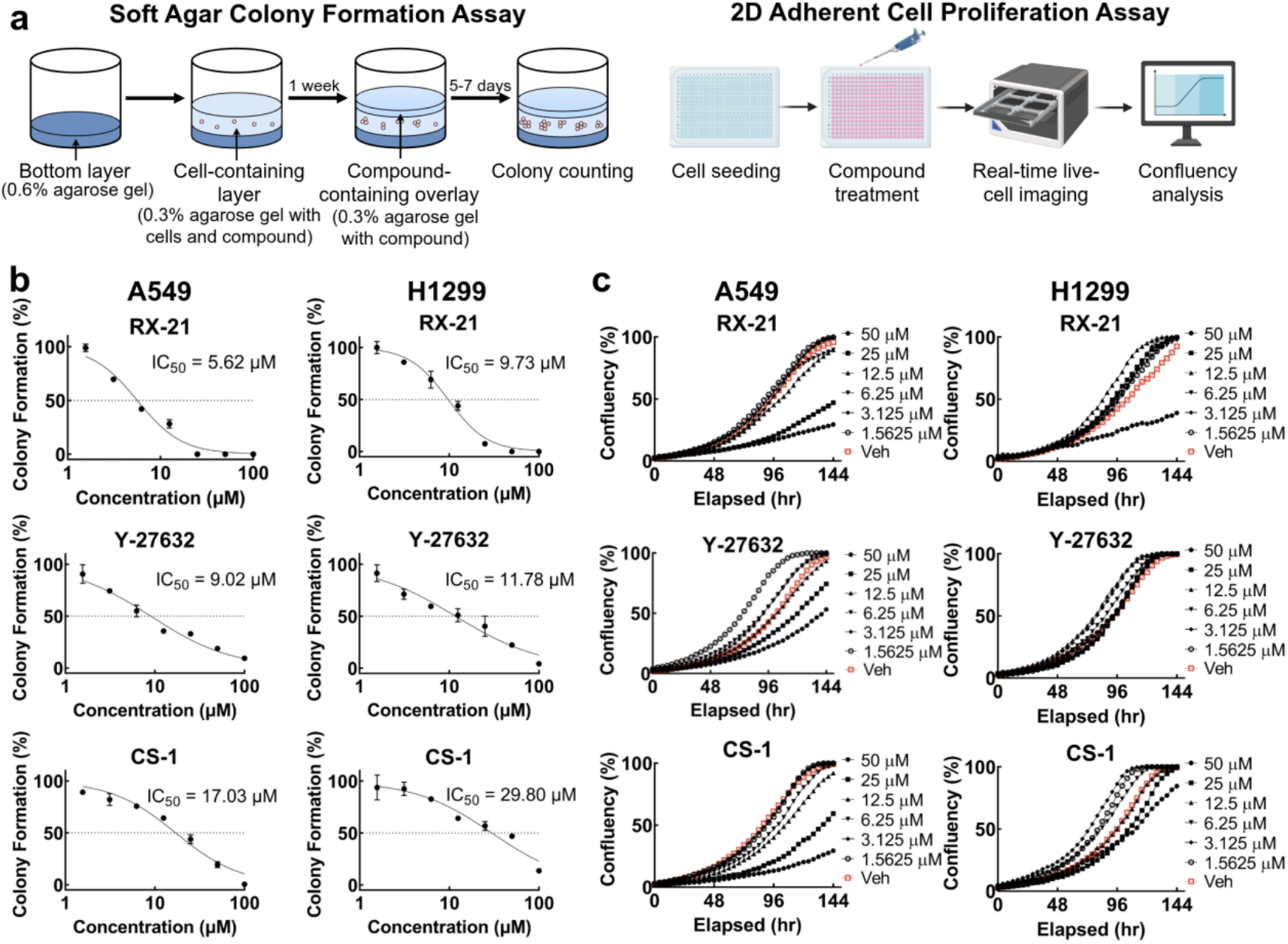
Effects of ROCK1 inhibitors on anchorage-independent growth and 2D proliferation in cancer cell lines. (a) Schematic of the experimental workflow. (b,c) Effects of RX-21, Y-27632 and CS-1 on anchorage-independent colony formation (b) and 2D proliferation (c) in A549 and H1299 cells. Soft agar colony formation was used as the primary readout of anchorage-independent growth, whereas the 2D proliferation assay was used to assess whether colony suppression was accompanied by broader inhibition of adherent cell growth. Colony formation data are presented as mean ± s.d. from three independent replicates. Dose-response curves were fitted to all replicate measurements using a variable-slope nonlinear regression model in GraphPad Prism, and the corresponding best-fit IC50 values are indicated in each panel. The dashed line marks 50% colony formation. For the 2D proliferation assay, two technical replicates were measured at each concentration, confluency was normalized to the vehicle control, with the maximum vehicle signal set to 100%.

## 3. CONCLUSION AND DISCUSSIONS

In this work, we present REAL-SWIT as a generative framework for ultra-large virtual screening within make-on-demand chemical space. The central premise is that a combinatorial library such as REAL Space should not be treated only as an enormous set of enumerated molecules, but also as a learnable distribution defined by building blocks, reaction types, and their allowable combinations. By learning this distribution from representative subsets, the resulting generator achieved approximately 96% REAL Space consistency, showing that generative sampling can remain largely within the underlying synthesizable library. Coupled with target-specific docking-surrogate guidance, REAL-SWIT further enables direct generation and prioritization of complete, synthesizable molecules without exhaustive enumeration or stepwise fragment expansion. Computational benchmarks and analyses, together with experimental ROCK1 validation, establish REAL-SWIT as a practical ULVS strategy that links the scale of make-on-demand libraries with the efficiency and flexibility of molecular generation.

A key observation from this work is that the learnability of REAL Space follows a clear data-scaling trend. As the query-based training set increased from 1 M to 4 M to 23 M, REAL Space consistency improved progressively, indicating that larger and more representative subsets enable the generator to better approximate the underlying distribution of the combinatorial library. This suggests that, in principle, further scaling of training data should continue to improve in-library consistency and component-level coverage, although with diminishing returns expected as the dominant BB and reaction-type patterns become saturated. Importantly, the improvement was not determined by training-set size alone. The query-based strategy produced a more balanced BB-frequency distribution than random sampling, allowing lower-frequency BBs to be represented sufficiently often for the model to learn their usage patterns. Thus, learning a combinatorial chemical space requires not only broad molecular coverage but also sufficient repeated exposure to the component-level grammar of the library.

The evaluation against later REAL Space releases further supports this interpretation. Molecules generated by models trained on earlier REAL Space releases were later found in expanded releases, showing that the generator did not simply memorize molecules from a fixed enumerated set. Instead, it learned structural and compositional rules that remained compatible with future expansions of REAL Space. In this sense, the model partially anticipated molecules that were not available in the training release but became available as the library evolved. This transferability is especially encouraging because it suggests that the model has learned chemical patterns and compositional rules underlying the construction and expansion of make-on-demand libraries. REAL Space represents only one implementation of this broader concept, and the same strategy should in principle be extendable to other make-on-demand libraries. Such an extension represents a natural and ongoing direction for broadening generative ULVS across diverse synthesizable chemical spaces.

REAL-SWIT also provides a distinct perspective on how ultra-large combinatorial spaces can be searched. Fragment- and synthon-based methods such as V-SYNTHES, Chemical Space Docking, and SyntheMol reduce the search space by prioritizing building blocks or capped fragments before expanding them into complete molecules. This strategy is highly efficient, but it necessarily relies on chemically incomplete or modified intermediates as proxies for the final assembled compounds. BBs often contain reactive handles, and capped fragments or synthons are only approximations of their assembled chemical states. Our results, together with recent observations from V-SYNTHES2, highlight that this approximation can become limiting when fragment poses, fragment scores, or BB contributions depend strongly on the final molecular context^24^. By generating and prioritizing complete REAL Space molecules directly, REAL-SWIT can capture context-dependent and cooperative BB combinations that are less readily accessible to fragment-level prioritization.

REAL-SWIT remains computationally efficient while shifting the primary search unit from isolated components to full molecules. Docking remains the principal computational bottleneck, because it is required to generate surrogate-training data and to validate top-ranked molecules, but REAL-SWIT reduces the need for exhaustive docking by concentrating docking calculations on selected generated subsets. In our implementation, CB2 screening with ICM-Pro could be completed in less than four days using a computational node equipped with a single A40 GPU and 72 CPU cores, with docking and generation accounting for about 60% and 25% of the total runtime, respectively (Table S2). Increasing the docking budget expanded the surrogate-training set and improved molecular diversity, while keeping the total runtime within a practical range (Figure S8).

The ROCK1 experiments provide a direct test of whether generated molecules can move beyond computational prioritization to experimentally measurable activity. From 23 synthesized compounds, six showed biochemical inhibition of ROCK1, including RX-3 with a submicromolar IC50 of 0.17 µM and multiple additional low-micromolar hits. These compounds were not close analogs of reported ROCK1 inhibitors, supporting the ability of REAL-SWIT to identify novel, synthesizable hits from REAL Space.

At the same time, the validation results illustrate a broader reality of AI- and physics-based VS: experimental readouts are indispensable, but they are context-dependent measurements rather than universal labels of compound quality. Apparent biochemical potency such as IC_50_ values can depend strongly on assay format and conditions, including ATP concentration for ATP-competitive kinase inhibitors, which may contribute to differences between our measurements and previously reported values. Cellular assays introduce additional layers of complexity, including compound permeability, intracellular stability, and phenotype-specific responses. Consistent with this complexity, RX-3 was the strongest biochemical inhibitor, whereas RX-21 showed the most favorable cellular activity among the compounds tested. These observations highlight that experimental validation is better viewed as a sequence of complementary filters in hit discovery rather than as a single definitive test for algorithmic success.

A remaining limitation of the current implementation is that REAL-SWIT is guided primarily by an empirical docking objective. This choice is practical and effective for large-scale enrichment, but docking scores are imperfect and can encode systematic preferences, for example biasing towards larger molecules with more aromatic rings^1,35^. As shown by the molecular-weight shift in the generated molecules, an efficient generative model can amplify such features when they are aligned with the optimization objective. Thus, while REAL-SWIT largely alleviates the synthesizability problem by operating within REAL Space, it also makes the quality of the objective function more central. In this sense, the next challenge for generative ULVS is not only to generate synthesizable molecules, but also to define optimization targets that better reflect the desired pharmacological profiles.

This perspective suggests several natural extensions. At the generation stage, target-specific surrogate models could be trained or fine-tuned using experimental data, such as high-throughput screening (HTS) or DNA-encoded library (DEL) results, when available. Alternatively, surrogate models could be constructed by distillation from co-folding-based affinity-prediction models such as Boltz-2^48^. In parallel, REAL-SWIT can be extended from single-objective to multi-objective generation, incorporating cLogP, solubility, and predicted ADMET properties. After the generative search has reduced the candidate space to a smaller prioritized set, simulation-based methods, in particular absolute or relative binding free energy calculations, could be applied for rescoring. More broadly, REAL-SWIT points toward a workflow in which make-on-demand libraries provide the synthesizable substrate, generative models enable efficient exploration, and increasingly rigorous objective functions determine which generated molecules are advanced toward experiment.

## Data and Code Availability

All source code, datasets, and model checkpoints required to reproduce the results reported in this study are publicly available at https://github.com/JingHuangLab/REAL-SWIT. The enumerated REAL compounds in ZINC-22 and the compounds in the ChEMBL dataset were obtained from https://cartblanche.docking.org/search/random and https://chembl.gitbook.io/chembl-interface-documentation/downloads, respectively. The template-defined REAL Space a datasets corresponding to the 36B, 76B, and 83B releases are available from the BioSolveIT Chemical Spaces resource at https://www.biosolveit.de/chemical-spaces.

## Supporting Information Available

Architectures and training curves of the molecular generative models (Figures S1 and S2); Maximum Tanimoto similarity of generated molecules absent from REAL Space to REAL Space molecules (Figure S3); analyses of building block and reaction coverage for the QB-L and RS models (Figures S4); comparison of ICM docking results for active molecules (Figure S5); comparison of docked poses of REAL-SWIT molecules, constituent fragments, and SyntheMol molecules (Figure S6); anchorage-independent colony formation and 2D adherent proliferation assays for five generated compounds and one reference compound (Figure S7); effects of minimum docking dataset size on molecular diversity and score distributions (Figure S8); REAL Space consistency of generated molecules across REAL Space releases (Table S1); computational details, including the time and resources required for REAL-SWIT (Table S2); REAL Space active compounds sharing building blocks with generated molecules and identification of better-scoring BB-sharing molecules (Table S3); and structures of ROCK1 compounds selected for experimental testing (Table S4).

## Acknowledgments

This work is supported by the Shenzhen Medical Research Fund (B2404003), the National Natural Science Foundation of China (T2596084), the Yangtze River Delta Joint Sci-Tech Innovation and Research Projects (2023CSJZN0600), the Westlake University-Muyuan Joint Research Institute (WU2025MY001), the State Key Laboratory of Gene Expression, and the Westlake Education Foundation. We thank Feng Xu from the Westlake University High-Throughput Core Facility for assistance with cell lines, the Westlake University Supercomputer Center for computational resources and related assistance, and the Enamine team for their valuable data support.

## MATERIALS AND METHODS

### Construction of representative subsets of REAL Space

Two types of datasets were constructed for training the molecular generative model. Query-based subsets were curated from REAL Space using SpaceLight v1.5.0^20^, the command-line implementation of the infinisee similarity-search suite^49^, with ECFP4 fingerprints. Specifically, each compound in ChEMBL v33 (∼2.3 million compounds) was used as a query in SpaceLight to retrieve the top k most similar compounds from the March 2023 release of REAL Space (∼36 billion compounds)^50^. To generate training sets at different scales, k was set to 1, 100, or 1,000. After deduplication, these searches yielded 1.5 million, 114 million, and 857 million unique compounds, respectively. The 1.5-million-compound set was used directly as the smallest query-based dataset, QB-S.

To construct medium- and large-scale query-based datasets that were both diverse and computationally tractable, the 114-million- and 857-million-compound sets were further reduced by clustering. Specifically, the two sets were partitioned into 1,140 and 8,570 clusters, respectively, using K-means clustering implemented in Faiss^51^, corresponding to approximately 100,000 compounds per cluster. Representative molecules were then selected from each cluster using a pairwise similarity cutoff of 0.45, yielding final datasets of 4 million and 23 million compounds, denoted QB-M and QB-L, respectively.

As a random-sampling baseline, 51 million molecules were randomly selected from ZINC-22 using its “Select random molecules” function^11^. Because ZINC-22 contains molecules from multiple make-on-demand libraries, including Enamine REAL Database, Enamine REAL Space, WuXi, and Mcule, SpaceLight was used to identify molecules originating from REAL Space. As the current ZINC-22 collection contains approximately 97 billion 2D molecules, substantially more than were available when ZINC-22 was originally reported, we screened the sampled molecules against two REAL Space releases to account for REAL Space molecules added to ZINC-22 after its initial publication: the March 2023 release (∼36 billion compounds) and the March 2025 release (∼76 billion compounds). Molecules assigned a similarity score of 1 in either search and canonical SMILES identical to the corresponding matched REAL Space molecule were considered present in REAL Space. This procedure yielded a 24-million-compound random-sampling baseline dataset.

Both the query-based and random-sampling datasets were subsequently preprocessed for model training. Molecules were removed if they contained fewer than 6 or more than 70 heavy atoms, more than 10 rings, a largest ring more than 8 atoms, or an aliphatic carbon chain longer than 4 atoms. Additional filters excluded molecules with a carbon atom fraction below 0.5, more than 91 tokens, a token-to-atom ratio greater than 2, or any token occurring at a frequency below 0.05. For each dataset, 100,000 molecules were randomly reserved for validation, and the remaining molecules were used for training. After preprocessing and validation-set removal, the training sets contained 1,450,835 molecules for QB-S, 4,116,646 for QB-M, and 23,381,021 for QB-L. For a direct comparison with QB-L, the RS training set was subsampled to the same size, yielding 23,381,021 molecules.

### Architecture and training of the generative model

During training, the SMILES strings of the input molecules were tokenized and encoded as vocabulary indices, which were then transformed into 1024-dimensional vectors through an embedding layer. The resulting embeddings were processed by six stacked LSTM layers, each with a hidden-state dimension of 2048. The output of the final LSTM layer was then projected through a linear layer onto the full token vocabulary, followed by a Softmax function to obtain the probability distribution over the next token (Figure S1). Training was performed using teacher forcing, in which the ground-truth token at each step was provided as the input for predicting the subsequent token. Model parameters were optimized by minimizing the negative log-likelihood of each molecular sequence, calculated as the sum of the negative log-probabilities assigned to the ground-truth tokens at all sequence positions:

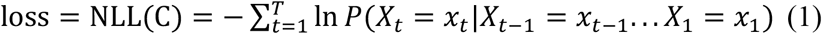

Where *P*(*X_t_* = *x_t_*|*X_t_*_−1_ = *x_t_*_−1_… *X*_1_ = *x*_1_) represents the conditional probability of token *x_t_* at step *t*, given the sequence of preceding tokens. Model training was performed using the Adam optimizer with a batch size of 512 for up to 300 epochs. The initial learning rate was set to 1 × 10^−4^, and was reduced by a factor of 0.8 if the validation loss did not improve for eight consecutive epochs. Training was terminated when the learning rate decreased to 1 × 10^−5^. The model with the lowest average negative log-likelihood on the validation set was selected for downstream applications (Figure S2). During sampling, generation was initiated with a start token, and subsequent tokens were sampled sequentially from the model-predicted probability distribution conditioned on the previously generated sequence until an end token was produced.

### Evaluation metrics

To assess the sampling performance of the trained generative models, each model was sampled in five independent runs of 5 million compounds each. The resulting molecular sets were evaluated in terms of validity, uniqueness, internal diversity, novelty, REAL Space consistency, building-block coverage, and reaction-type coverage. Validity was defined as the fraction of sampled SMILES strings that could be successfully parsed by RDKit. Uniqueness was defined as the fraction of distinct canonical SMILES among valid sampled molecules. Internal diversity was calculated following the MOSES benchmark from the average pairwise Tanimoto similarity among generated molecules, with lower similarity corresponding to higher diversity^52^. Novelty was defined as the fraction of valid sampled molecules not present in the corresponding training set. REAL Space consistency was defined as the fraction of sampled molecules identified by SpaceLight as present in the REAL Space, considering the 36B, 76B, and 83B releases. Molecules matched to REAL Space were further analyzed for chemical space coverage by counting the number of unique building blocks and reaction types represented in the sampled molecules. For each matched molecule, the reaction and reagent names reported by SpaceLight were parsed. Reagent names were mapped to the corresponding building blocks using information provided by Enamine (October 2023), and the resulting assignments were used to calculate building-block coverage. Reaction coverage was calculated using the major reaction classes indicated by the numerical identifiers in the SpaceLight reaction names, without distinguishing subclasses denoted by additional letters.

### Architecture and training of the docking surrogate model

For each target protein, compounds with associated docking scores were randomly split into training and test sets at a 9:1 ratio. Molecules were represented as graphs, with atoms as nodes and bonds as edges, and the resulting graph representations served as inputs to a Directed Message Passing Neural Network (D-MPNN) implemented in the Chemprop framework^53,54^. During message passing, 300-dimensional hidden states associated with directed bonds were iteratively updated over three message-passing steps. In the readout stage, incoming bond messages were aggregated for each atom and combined with the corresponding atom features to obtain atom-level representations, which were then mean pooled to generate a molecular-level embedding. This embedding was then passed through two fully connected layers, each with 300 hidden units and ReLU activation, to predict the target-specific docking score. Model parameters were optimized by minimizing the L2 loss between predicted and observed docking scores using the Adam optimizer. Training was performed for up to 50 epochs with a batch size of 50.

### Target-guided molecular generation with REAL-SWIT

Target-guided molecular generation was performed within the REINVENT 2.0 reinforcement-learning framework^37^. The REAL Space generative model trained in this study was used as both the prior and initial agent, while the target-specific D-MPNN docking surrogate model provided the target-directed scoring component. This setup was designed to bias generation toward synthesizable chemical space while favoring molecules with improved predicted docking scores.

The molecular scoring function used in this study comprised two components: target-specific score and molecular weight. The framework also allows additional scoring components to be incorporated for other generation objectives. To place the two scoring terms on a common scale for their combination in the multi-parameter objective, each was transformed to a value between 0 and 1. The transformation parameters were predefined to specify the desired score ranges and transition profiles. For each generated molecule, the D-MPNN-predicted target-specific score d was transformed using a sigmoid function, with more favorable (more negative) scores mapped closer to 1, and *a*_2_ = 1,

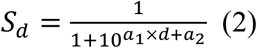

where *a_1_ = 1/30 and a_2_ = 1*. Molecular weight was also transformed to a 0-1 score, with molecules between 200 and 600 Da mapped closer to 1 and those outside this range progressively penalized,

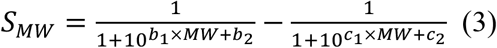

where *b*_1_ = *c*_1_ = − 1/30, *b*_2_ = 20/3, and *c*_2_ = 20. The transformed target-specific and molecular-weight scores were combined as a weighted sum to define the multi-parameter objective (MPO) score, subject to the REINVENT IdenticalTopologicalScaffold diversity filter, which set the MPO score to 0 when the number of molecules sharing the same topological Murcko scaffold exceeded a predefined limit,

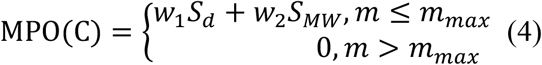

where the weighting factors were set to *w*_1_ = 2/3 and *w*_2_ = 1/3; m denotes the number of molecules in memory sharing the same topological Murcko scaffold as molecule C; and *m_max_* denotes the maximum allowed number per scaffold and was set to 10 in this study.

The MPO score was then combined with the prior negative log-likelihood to define the augmented negative log-likelihood, which served as the optimization objective for the agent by favoring high-scoring molecules while retaining the molecular distribution learned by the prior,

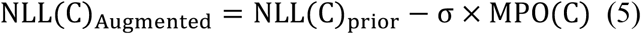

where *σ* was set to 128. The agent was optimized to match this objective by minimizing the squared difference between the agent and augmented negative log-likelihoods,

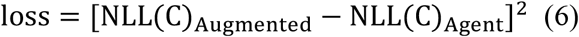

using the Adam optimizer.

For each protein target, reinforcement learning was performed for up to 15,000 steps with a batch size of 128. At each step, the agent generated a batch of molecules, whose MPO scores were used to calculate the augmented negative log-likelihoods and guide the parameter update according to Eq. (6). Generated molecules were accumulated in memory throughout the process, which was terminated once the memory contained more than 1 million molecules. Finally, the resulting molecules were ranked by target-specific score, and the top 250,000 were screened against REAL Space using SpaceLight. Among molecules identified as present in the designated REAL Space releases, the top 100,000 were selected for subsequent docking calculations.

### Molecular docking and similarity calculations

To enable direct comparison with V-SYNTHES, we evaluated the same two protein targets, human cannabinoid receptor 2 (CB2; PDB ID: 5ZTY) and Rho-associated protein kinase 1 (ROCK1; PDB ID: 2ETR)^55,56^, and performed protein-ligand docking using ICM-Pro (version 3.8-7b)^38^. Receptor preparation closely followed the procedure described for V-SYNTHES. Receptors were prepared by converting the PDB structures into ICM internal coordinate objects, during which missing heavy atoms and hydrogen atoms were added, polar hydrogens were optimized, and the protonation states and side-chain rotamers of His, Asn, and Gln residues were adjusted. The binding sites were defined by residues surrounding the crystallographic ligands, comprising 30 residues for CB2 and 20 residues for ROCK1, and the corresponding docking maps were subsequently generated. Docking was then performed in batch mode using the _dockScan function with an effort value of 2, generating up to 16 poses per ligand. Each ligand was docked in two independent runs, and the more favorable of the two docking scores was used as the final score for each ligand-target pair.

For structural comparison of molecules, 2D similarity was calculated as the Tanimoto similarity between ECFP4 fingerprints generated with RDKit, whereas 3D similarity was assessed by rigid-body molecular alignment using LS-align42.

### Implementation of SyntheMol

SyntheMol was run using the publicly available code from its GitHub repository. To ensure a consistent comparison with REAL-SWIT, the building blocks supplied with SyntheMol were docked under the same conditions used for complete molecules, and the resulting scores were used as precomputed building-block scores. The same CB2-specific and ROCK1-specific D-MPNN models used in REAL-SWIT were also employed for target-guided scoring. For each target, 1,000,000 rollouts were performed, after which the generated molecules were ranked by predicted score and the top 100,000 were selected for docking and subsequent comparison of screening performance.

### Selection of ROCK1 candidates for experimental validation

A total of 100,000 ROCK1-targeted molecules generated by REAL-SWIT were subjected to a multi-step filtering and prioritization workflow. Molecules violating more than one of Lipinski’s rules or containing PAINS motifs, identified using the RDKit FilterCatalog, were removed, leaving 96,401 molecules. To enrich for compounds capable of engaging the ATP-binding site, the remaining molecules were further filtered by requiring a hydrogen-bonding interaction with the hinge residue Met156. This constraint was applied using the pharmacophore query function in MOE under the Extended Hückel Theory (EHT) scheme (R = 1.6, average sphere radius; S = Acc > 1, minimum required strength of a matching ligand atom)^57^, yielding 54,733 candidates.

To reduce redundancy while retaining diverse protein-ligand interaction patterns, the retained docking poses were analyzed using ProLIF to generate protein-ligand interaction fingerprints^43^. Each molecule was represented as a 115-bit binary vector encoding the presence or absence of specific interaction types. The fingerprints were then clustered by agglomerative clustering implemented in scikit-learn (average linkage; distance threshold = 0.38), resulting in 1,416 clusters. From each cluster, the highest-ranked molecule was retained, with additional high-scoring molecules selected from larger clusters, yielding 2,123 molecules for manual inspection. Visual inspection then selected 28 compounds with distinct hinge-binding core scaffolds, 23 of which were successfully procured from Enamine at >90% purity for experimental validation.

### ROCK1 enzymatic activity assay

ROCK1 enzymatic activity was measured using a luminescent ADP-Glo^TM^ kinase assay (Promega, Cat. #V9101), which quantifies ADP generated during the kinase reaction. Recombinant human ROCK1 (Promega, Cat. #V3411) was used in all experiments. GSK429286A (TopScience, Cat. #T2633) was used as a positive control, whereas RS-15 (Enamine, Cat. #Z1006542468) and CS-1 (Enamine, Cat. #Z4540472702), reported in previous ULVS studies, were included as reference compounds. Compounds were tested in technical duplicate, and each experiment was independently repeated three times to determine inhibition values and IC_50_ values. The final assay concentrations were 9.5 nM ROCK1, 10 µM ATP, and 50 µM S6K peptide substrate (KRRRLASLR). Assays were performed in white 384-well plates (PerkinElmer, Cat. #6007290). Briefly, 19 µL of ROCK1 (20 nM) prepared in reaction buffer (25 mM HEPES, pH 7.4; 150 mM NaCl; 20 mM MgCl2; 0.01% Triton X-100; 0.01% BSA; 5 mM DTT) was mixed with 1 µL of serially diluted compound and incubated at room temperature for 40 min. The kinase reaction was initiated by adding 20 µL of a substrate mixture containing ATP (20 µM) and S6K peptide (100 µM) in reaction buffer. Plates were then incubated at 37 °C for 60 min. Following the kinase reaction, 10 µL of each reaction mixture was transferred to a new 384-well plate for detection. The ADP-Glo^TM^ detection was performed according to the manufacturer’s instructions in two steps. First, 10 µL of ADP-Glo^TM^ Reagent was added to terminate the kinase reaction and deplete remaining ATP. Second, 20 µL of Detection Reagent was added to convert ADP to ATP and enable luminescence detection via a luciferase/luciferin reaction. Luminescence was measured using a plate luminometer. Control reactions without ROCK1 were included to determine background signal. After subtraction of background values, data were normalized to reactions containing DMSO alone. IC_50_ values were calculated using GraphPad Prism.

### Cell culture

A549 cells were maintained in Dulbecco’s Modified Eagle Medium (DMEM), whereas NCI-H1299 cells were maintained in RPMI 1640 medium. Both media were supplemented with 10% FBS, 1× penicillin–streptomycin, and 2 mM L-glutamine. All cell lines were cultured at 37°C in a humidified incubator with 5% CO_2_ and were routinely tested for mycoplasma contamination.

### Anchorage-independent colony formation assay

Soft agar colony formation assays were performed to assess anchorage-independent cell growth in response to ROCK1 inhibition, as previously described with modifications^58^. Briefly, 12-well plates were coated with 700 μL of pre-warmed 0.6% low-gelling agarose (Sigma-Aldrich) prepared in basal cell culture medium and allowed to solidify at room temperature to form the base layer. ROCK1 inhibitors or vehicle control (DMSO) were serially diluted and mixed with 1 × 10³ cells in 0.6% low-gelling agarose supplemented with 10% FBS to achieve a final agarose concentration of 0.3%. The cell-agarose mixture was overlaid onto the base layer and allowed to solidify at room temperature. Pre-warmed compound dilution in 0.3% agarose was supplemented 7 days post-seeding. Colonies consisting of more than 10 cells were counted in five representative fields per well at 12–14 days post-seeding. All experiments were performed in triplicates, and data are presented as relative colony formation compared with the vehicle control (mean ± SD).

### 2D adherent proliferation assay

A549 and NCI-H1299 cells were seeded at a density of 500 cells per well in 384-well plates and allowed to adhere for 24 h. The culture medium was then replaced with fresh medium containing serial dilutions of the indicated compounds or vehicle control, with two technical replicate wells per condition. Cell growth was monitored using an Incucyte SX5 live-cell imaging system (Sartorius) and maintained under standard culture conditions (37 °C, 5% CO_2_). Images were acquired every 4 h using a 10× objective over a period of 6 days. Cell confluency was quantified using Incucyte SX5 software and plotted as a function of time.

## REFERENCE

1. Lyu, J. et al. Ultra-large library docking for discovering new chemotypes. Nature 566, 224–229 (2019).

2. Liu, F. et al. The impact of library size and scale of testing on virtual screening. Nat. Chem. Biol. 21, 1039–1045 (2025).

3. Stein, R. M. et al. Virtual discovery of melatonin receptor ligands to modulate circadian rhythms. Nature 579, 609–614 (2020).

4. Alon, A. et al. Structures of the σ2 receptor enable docking for bioactive ligand discovery. Nature 600, 759–764 (2021).

5. Fink, E. A. et al. Structure-based discovery of nonopioid analgesics acting through the α2A-adrenergic receptor. Science 377, eabn7065 (2022).

6. Singh, I. et al. Structure-based discovery of conformationally selective inhibitors of the serotonin transporter. Cell 186, 2160–2175.e17 (2023).

7. Liu, F. et al. Structure-based discovery of CFTR potentiators and inhibitors. Cell 187, 3712–3725.e34 (2024).

8. Liu, F. et al. Large library docking identifies positive allosteric modulators of the calcium-sensing receptor. Science 385, eado1868 (2024).

9. Grygorenko, O. O. et al. Generating multibillion chemical space of readily accessible screening compounds. iScience 23, 101681 (2020).

10. Patel, H. et al. SAVI, in silico generation of billions of easily synthesizable compounds through expert-system type rules. Sci. Data 7, 384 (2020).

11. Tingle, B. I. et al. ZINC-22—a free multi-billion-scale database of tangible compounds for ligand discovery. J. Chem. Inf. Model. 63, 1166–1176 (2023).

12. Gorgulla, C. et al. An open-source drug discovery platform enables ultra-large virtual screens. Nature 580, 663–668 (2020).

13. Cecchini, D. et al. AI-enhanced adaptive virtual screening of large libraries for ligand discovery. Nat. Biotechnol. (2026).

14. Gentile, F. et al. Deep Docking: A deep learning platform for augmentation of structure based drug discovery. ACS Cent. Sci. 6, 939–949 (2020).

15. Graff, D. E., Shakhnovich, E. I. & Coley, C. W. Accelerating high-throughput virtual screening through molecular pool-based active learning. Chem. Sci. 12, 7866–7881 (2021).

16. Yang, Y. et al. Efficient exploration of chemical space with docking and deep learning. J. Chem. Theory Comput. 17, 7106– 7119 (2021).

17. Gentile, F. et al. Artificial intelligence-enabled virtual screening of ultra-large chemical libraries with deep docking. Nat. Protoc. 17, 672–697 (2022).

18. Rarey, M. & Stahl, M. Similarity searching in large combinatorial chemistry spaces. J. Comput. Aided Mol. Des. 15, 497–520 (2001).

19. Lessel, U., Wellenzohn, B., Lilienthal, M. & Claussen, H. Searching fragment spaces with feature trees. J. Chem. Inf. Model. 49, 270–279 (2009).

20. Bellmann, L., Penner, P. & Rarey, M. Topological similarity search in large combinatorial fragment spaces. J. Chem. Inf. Model. 61, 238–251 (2021).

21. Schmidt, R., Klein, R. & Rarey, M. Maximum common substructure searching in combinatorial make-on-demand compound spaces. J. Chem. Inf. Model. 62, 2133–2150 (2022).

22. Cheng, C. & Beroza, P. Shape-aware synthon search (SASS) for virtual screening of synthon-based chemical spaces. J. Chem. Inf. Model. 64, 1251–1260 (2024).

23. Sadybekov, A. A. et al. Synthon-based ligand discovery in virtual libraries of over 11 billion compounds. Nature 601, 452–459 (2022).

24. Nazarova, A. L. et al. V-SYNTHES2—the next generation tool for structure-based virtual screening of giga-scale chemical spaces. npj Drug Discov. 3, 21 (2026).

25. Beroza, P. et al. Chemical space docking enables large-scale structure-based virtual screening to discover ROCK1 kinase inhibitors. Nat. Commun. 13, 6447 (2022).

26. Eisenhuth, P., Liessmann, F., Moretti, R. & Meiler, J. Ultra-large library screening with an evolutionary algorithm in Rosetta (REvoLd). Commun. Chem. 8, 335 (2025).

27. Moesgaard, L. & Kongsted, J. Introducing SpaceGA: A search tool to accelerate large virtual screenings of combinatorial libraries. J. Chem. Inf. Model. 64, 8123–8130 (2024).

28. Swanson, K. et al. Generative AI for designing and validating easily synthesizable and structurally novel antibiotics. *Nat*. Mach. Intell. 6, 338–353 (2024).

29. Wang, M. et al. ClickGen: Directed exploration of synthesizable chemical space via modular reactions and reinforcement learning. Nat. Commun. 15, 10127 (2024).

30. Zhang, J. et al. Combinatorial docking and molecular generation to navigate over 100-billion molecules for prospective ligand discovery. Preprint at 10.64898/2026.06.07.730716 (2026).

31. Sadybekov, A. V. & Katritch, V. Computational approaches streamlining drug discovery. Nature 616, 673–685 (2023).

32. Barelier, S. et al. Substrate deconstruction and the nonadditivity of enzyme recognition. J. Am. Chem. Soc. 136, 7374–7382 (2014).

33. Peng, X., et al. Pocket2Mol: Efficient molecular sampling based on 3D protein pockets. In Proceedings of the 39th International Conference on Machine Learning 162, 17644–17655 (PMLR, 2022).

34. Cremer, J. et al. FLOWR: Flow matching for structure-aware de novo, interaction- and fragment-based ligand generation. Nat. Comput. Sci. 6, 565–574 (2026).

35. Zhang, W., Zhang, K. & Huang, J. A simple way to incorporate target structural information in molecular generative models. J. Chem. Inf. Model. 63, 3719–3730 (2023).

36. Gao, W., Luo, S. & Coley, C. W. Generative AI for navigating synthesizable chemical space. Proc. Natl Acad. Sci. USA 122, e2415665122 (2025).

37. Blaschke, T. et al. REINVENT 2.0: An AI tool for de novo drug design. J. Chem. Inf. Model. 60, 5918–5922 (2020).

38. Abagyan, R., Totrov, M. & Kuznetsov, D. ICM—a new method for protein modeling and design: Applications to docking and structure prediction from the distorted native conformation. J. Comput. Chem. 15, 488–506 (1994).

39. Zhang, W. & Huang, J. EViS: An enhanced virtual screening approach based on pocket-ligand similarity. J. Chem. Inf. Model. 62, 498–510 (2022).

40. Leeson, P. D. & Springthorpe, B. The influence of drug-like concepts on decision-making in medicinal chemistry. Nat. Rev. Drug Discov. 6, 881–890 (2007).

41. Ryckmans, T. et al. Rapid assessment of a novel series of selective CB(2) agonists using parallel synthesis protocols: A lipophilic efficiency (LipE) analysis. Bioorg. Med. Chem. Lett. 19, 4406–4409 (2009).

42. Hu, J., Liu, Z., Yu, D. J. & Zhang, Y. LS-align: An atom-level, flexible ligand structural alignment algorithm for high-throughput virtual screening. Bioinformatics 34, 2209–2218 (2018).

43. Bouysset, C. & Fiorucci, S. ProLIF: A library to encode molecular interactions as fingerprints. J. Cheminform. 13, 72 (2021).

44. Goodman, K. B. et al. Development of dihydropyridone indazole amides as selective Rho-kinase inhibitors. J. Med. Chem. 50, 6–9 (2007).

45. Vigil, D. et al. ROCK1 and ROCK2 are required for non-small cell lung cancer anchorage-independent growth and invasion. Cancer Res. 72, 5338–5347 (2012).

46. Watanabe, K. et al. A ROCK inhibitor permits survival of dissociated human embryonic stem cells. Nat. Biotechnol. 25, 681–686 (2007).

47. Sato, T. et al. Single Lgr5 stem cells build crypt-villus structures in vitro without a mesenchymal niche. Nature 459, 262–265 (2009).

48. Passaro, S. et al. Boltz-2: Towards accurate and efficient binding affinity prediction. Preprint at 10.1101/2025.06.14.659707 (2025).

49. Infinisee version 7.0.0, BioSolveIT GmbH, Sankt Augustin, Germany (2025).

50. Zdrazil, B. et al. The ChEMBL database in 2023: A drug discovery platform spanning multiple bioactivity data types and time periods. Nucleic Acids Res. 52, D1180–D1192 (2024).

51. Douze, M., et al. The Faiss library. IEEE Trans. Big Data 12, 346–361 (2026).

52. Polykovskiy, D. et al. Molecular sets (moses): A benchmarking platform for molecular generation models. Front. Pharmacol. 11, 565644 (2020).

53. Dai, H., Dai, B. & Song, L. Discriminative embeddings of latent variable models for structured data. In Proceedings of the 33rd International Conference on Machine Learning 48, 2702–2711 (PMLR, 2016).

54. Yang, K. et al. Analyzing learned molecular representations for property prediction. J. Chem. Inf. Model. 59, 3370–3388 (2019).

55. Li, X. et al. Crystal structure of the human cannabinoid receptor CB2. Cell 176, 459–467.e13 (2019).

56. Jacobs, M. et al. The structure of dimeric ROCK I reveals the mechanism for ligand selectivity. J. Biol. Chem. 281, 260–268 (2006).

57. Molecular Operating Environment (MOE), 2020.09; Chemical Computing Group ULC, Montreal, QC, Canada (2020).

58. Nakamura, D. The evaluation of tumorigenicity and characterization of colonies in a soft agar colony formation assay using polymerase chain reaction. Sci. Rep. 13, 5405 (2023).

