## Supplemental Figures and Tables for "Generative Access to Make-on-Demand Chemical Space Enables Ultra-Large Virtual Screening"

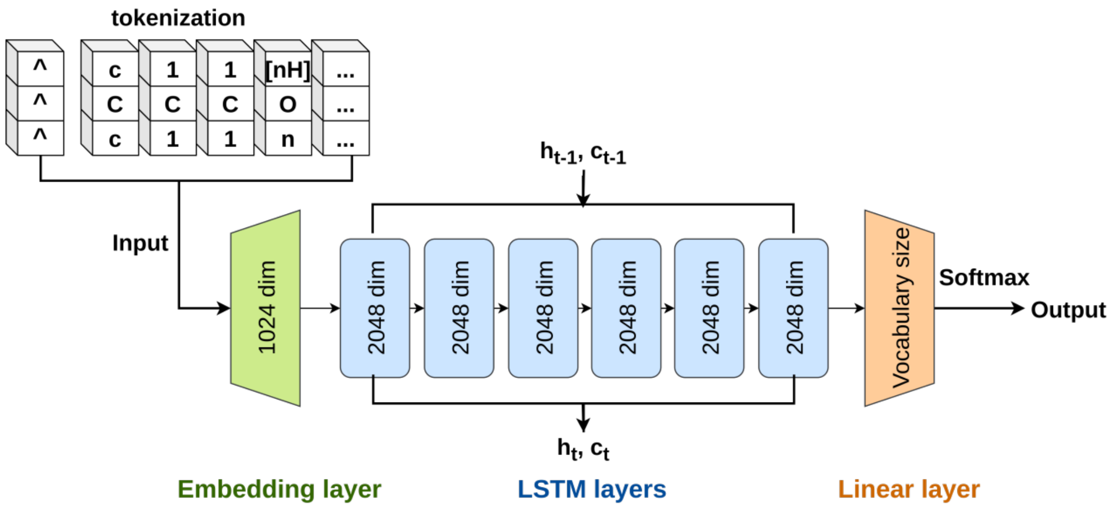


**Figure S1. Architecture of the LSTM-based molecular generative model used in REAL-SWIT.**

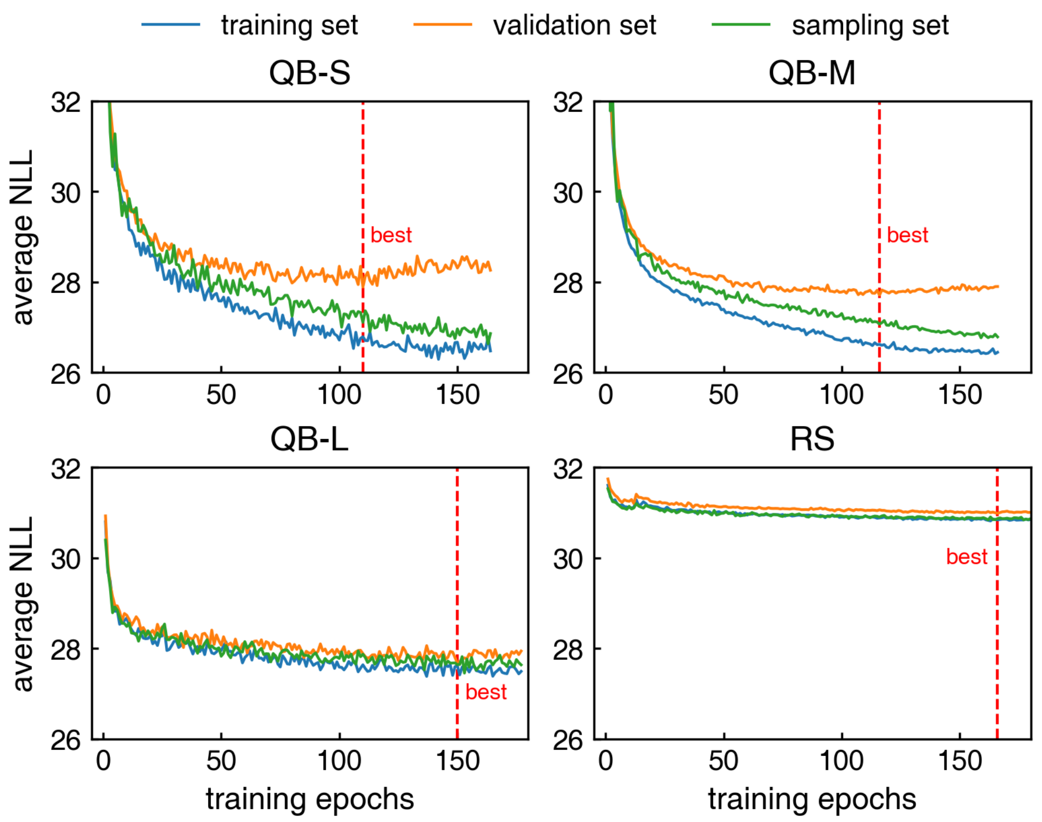


**Figure S2. Training curves of the four molecular generative models.** Blue, orange and green lines show the average negative log-likelihood (NLL) for the training, validation and sampling sets, respectively. For each model, the red dashed line indicates the epoch with the lowest validation NLL, which was selected as the model checkpoint.

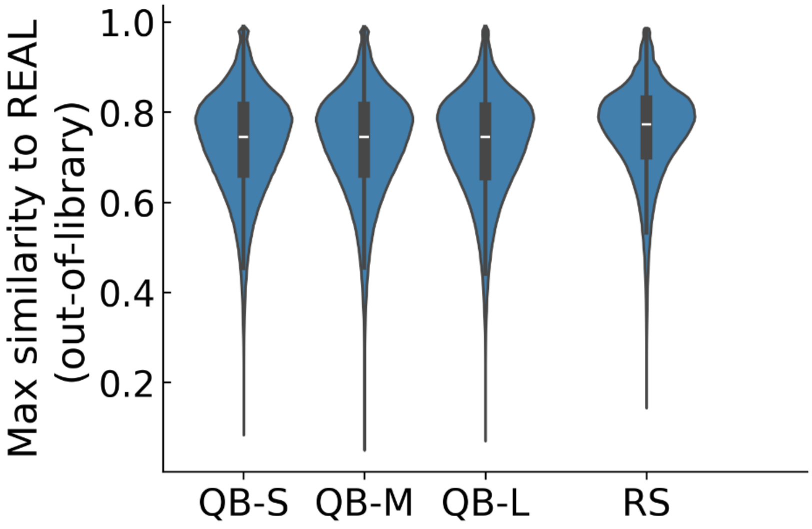


**Figure S3. Maximum Tanimoto similarity to REAL Space molecules for generated molecules absent from all examined REAL Space releases.** Distributions are aggregated across the five independent runs for each model.

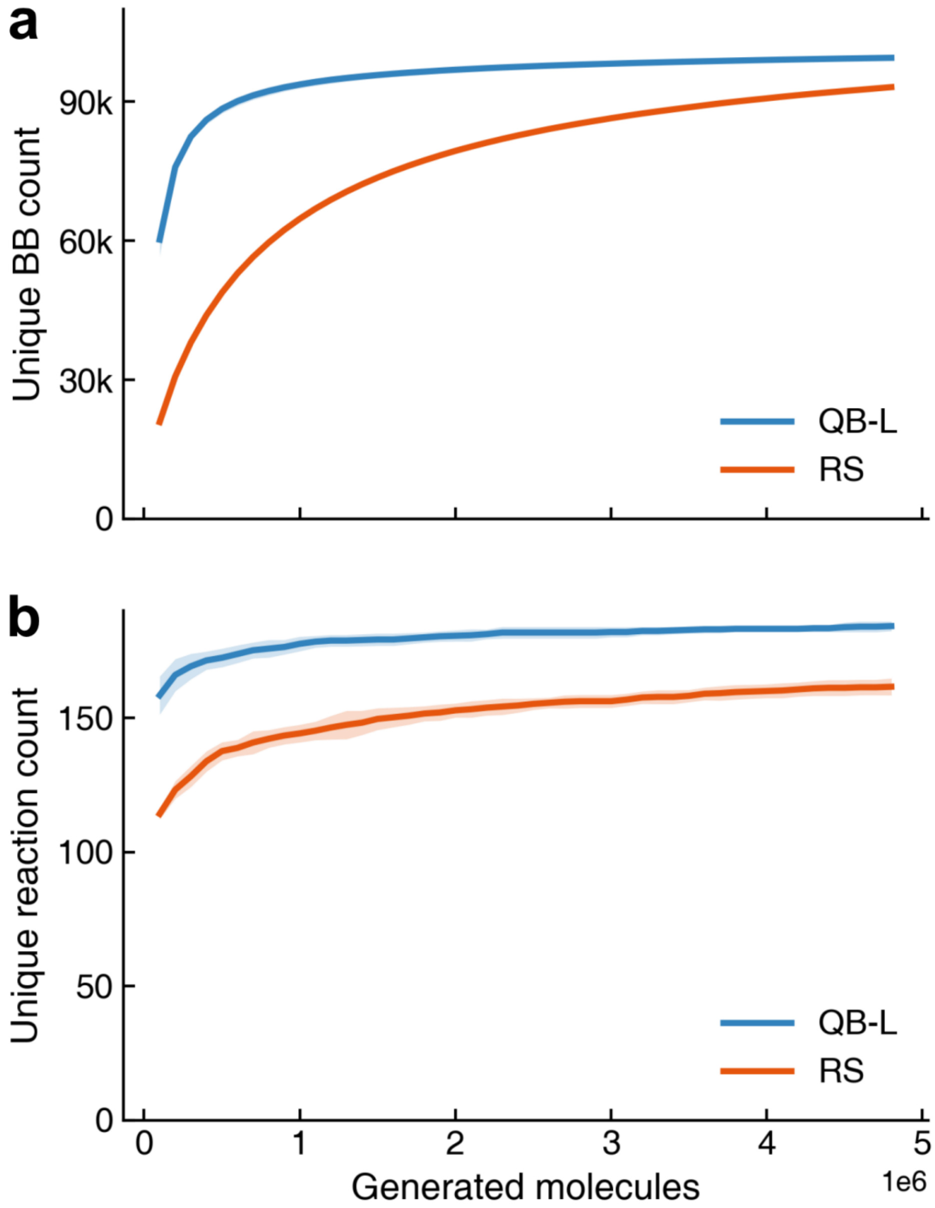


**Figure S4. Cumulative numbers of unique (a) building blocks and (b) reaction types recovered among molecules generated by the QB-L and RS models as the number of generated molecules increased.** Solid lines indicate the mean across five independent runs, and shaded regions represent mean ± s.d.

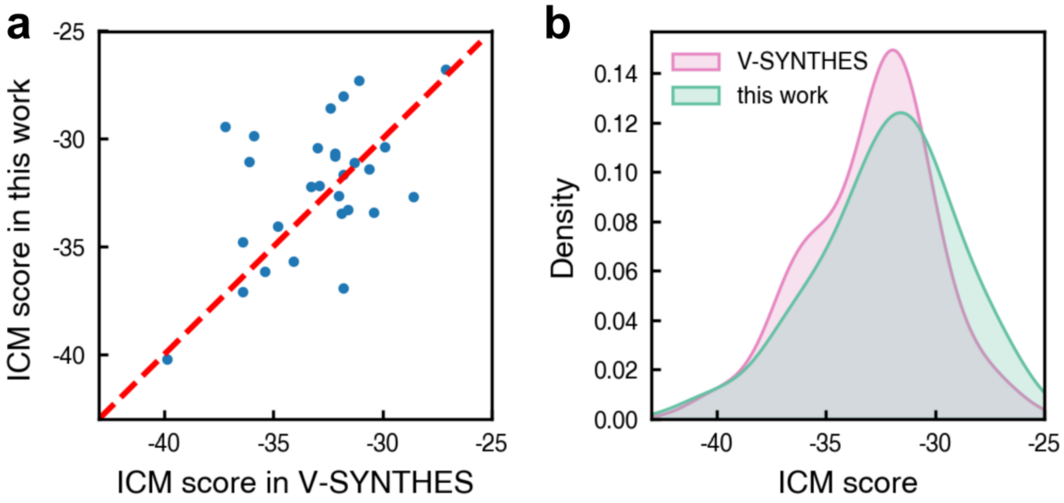


**Figure S5. Comparison of ICM docking scores for 28 active CB2 compounds reported in the V-SYNTHES study and obtained in this work.** (a) Scatter plot of the scores reported in the V-SYNTHES study against those obtained in this work. (b) Density distributions of the two sets of docking scores.

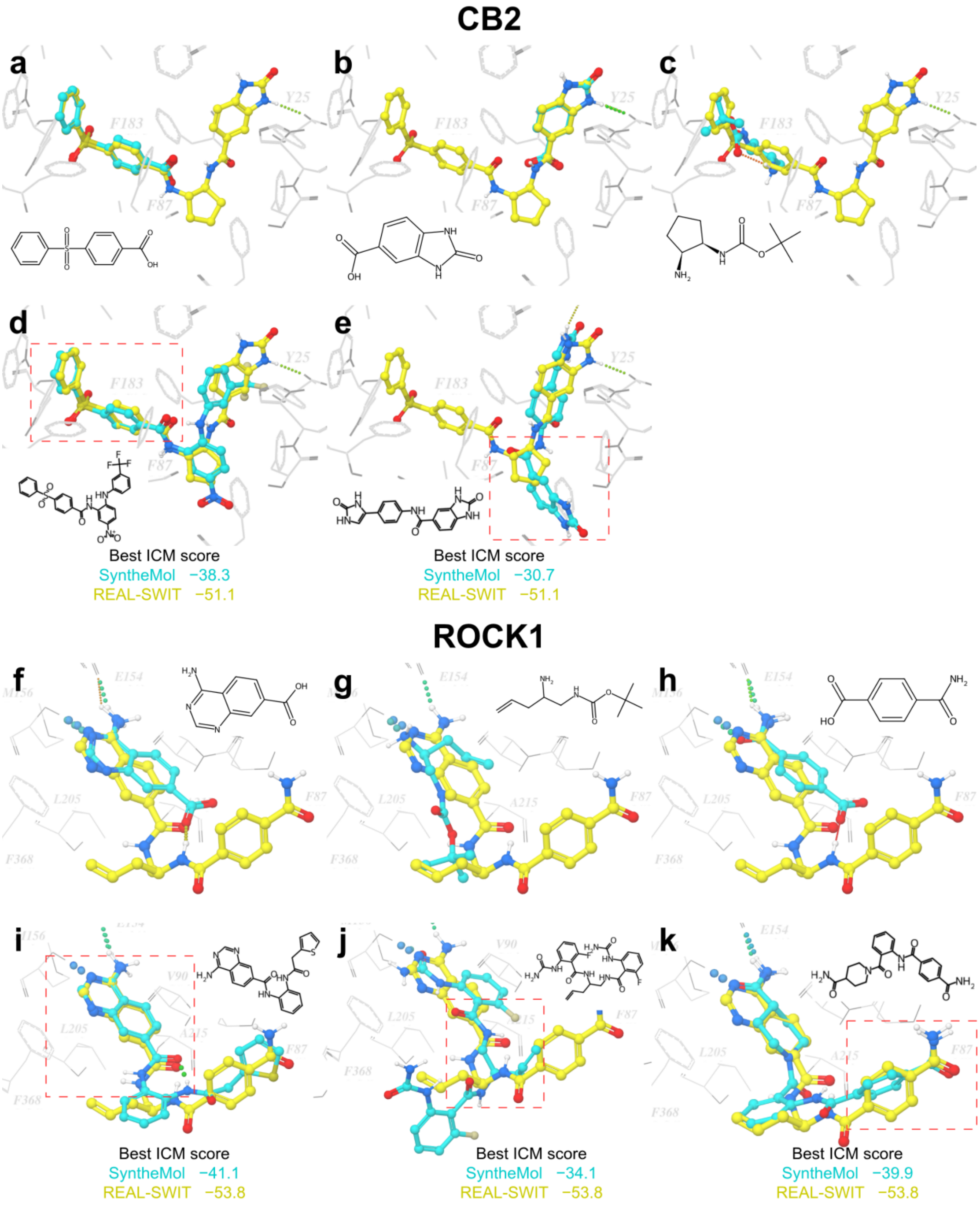


**Figure S6. Comparison of docked poses of representative REAL-SWIT molecules, their constituent fragments, and fragment-derived SyntheMol molecules for CB2 and ROCK1.** (a–c, f–h) Independently docked fragments (cyan) and the corresponding REAL-SWIT molecules (yellow) are shown in the CB2 and ROCK1 binding pockets, respectively. (d, e, i–k) The best-scoring SyntheMol molecule derived from the fragment shown directly above in the same column is shown in cyan together with the representative REAL-SWIT molecule in yellow. Two-dimensional structures of the cyan-colored fragments or SyntheMol molecules and the ICM docking scores of the compared full molecules are shown in each panel. Red dashed boxes indicate the positions of the corresponding fragments within the full molecules.

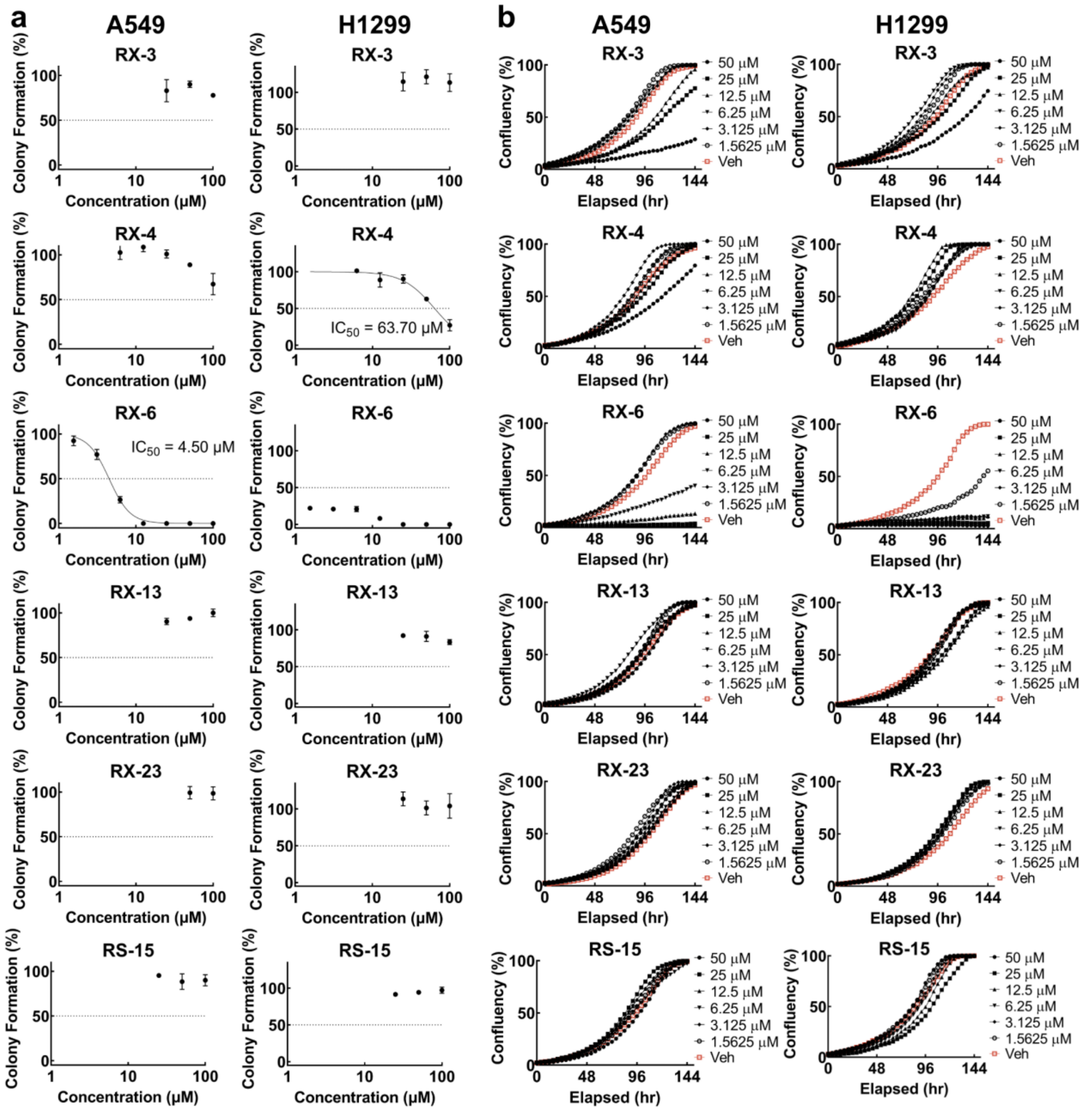


**Figure S7. Effects of five REAL-SWIT-generated ROCK1 hits and RS-15 on anchorage-independent colony formation and 2D adherent proliferation in A549 and H1299 cells.** (a) Anchorage-independent colony formation in A549 and H1299 cells treated with five REAL-SWIT-generated hits and RS-15, a ROCK1 hit identified by V-SYNTHES. (b) 2D adherent cell proliferation under the same compound treatments. Colony formation data are presented as mean ± s.d. from three independent replicates. Dose-response curves were fitted to all replicate measurements using a variable-slope nonlinear regression model in GraphPad Prism, and the corresponding best-fit $IC_{50}$ values are indicated where applicable. For the 2D proliferation assay, two technical replicates were measured at each concentration.

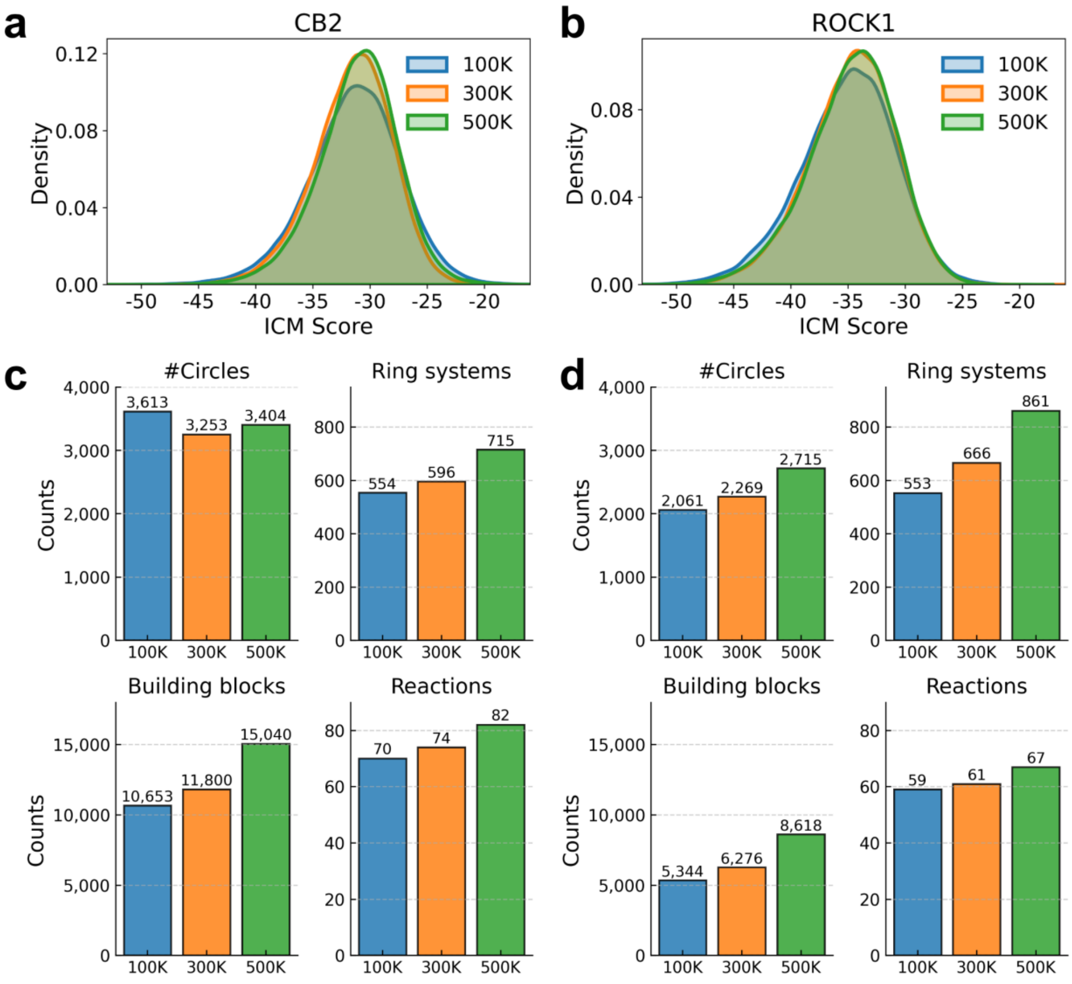


**Figure S8. Effect of docking dataset size on docking score distributions and structural diversity of generated molecules.** The 100K, 300K and 500K settings indicate that 100,000, 300,000 and 500,000 molecules, respectively, were docked in each of events 1 and 5 in Table S2, yielding approximately 0.2, 0.6 and 1.0 million docking results for the second round of D-MPNN training in event 6. For direct comparison, 100,000 molecules were docked in event 9 for all settings. (a, b) Docking score distributions for CB2- and ROCK1-generated molecules, respectively. (c, d) Scaffold diversity of CB2- and ROCK1-generated molecules, respectively. #Circles denotes the number of structurally distinct molecules at a Tanimoto similarity cutoff of 0.45. Ring systems denote the number of unique ring systems, with rings sharing at least one atom considered part of the same ring system. Building blocks and reactions were counted as described in Methods, Evaluation metrics. For CB2, the total REAL-SWIT runtimes for the 100K, 300K and 500K settings were 3 d 17 h, 6 d 8 h and 9 d 4 h, respectively.

**Table S1. REAL Space consistency of molecules generated by the four models across successive REAL Space releases.** Values are reported as mean ± s.d. across five independent runs, with 5 million molecules generated per run.

| Model | 36B release (%) | 76B increment (%) | 83B increment (%) | Overall REAL Space consistency (%) |
|---|---|---|---|---|
| QB-S | 83.20 ± 0.37 | 4.12 ± 0.29 | 0.41 ± 0.06 | 87.73 ± 0.03 |
| QB-M | 86.87 ± 0.08 | 3.70 ± 0.03 | 0.36 ± 0.02 | 90.93 ± 0.06 |
| QB-L | 94.65 ± 0.06 | 1.75 ± 0.13 | 0.139 ± 0.004 | 96.54 ± 0.07 |
| RS | 96.55 ± 0.17 (36B and 76B combined) | | 0.24 ± 0.13 | 96.79 ± 0.06 |

**Table S2. Computational steps, runtimes and resources of the REAL-SWIT workflow, using CB2 as an example, on NVIDIA A40 GPUs and Intel Xeon Gold 6150 CPUs (2.70 GHz).**

| Events | Runtime | Resources |
|---|---|---|
| 1. Docking of 100,000 sampled REAL compounds | 15 h 32 min | 72 CPU cores |
| 2. Initial D-MPNN training | 38 min | 1 GPU, 6 CPU cores |
| 3. Generation of the first 1,000,000 compounds | 10 h 45 min | 1 GPU, 6 CPU cores |
| 4. Identification of the top 100,000 REAL compounds generated in event 3 | 6 h 36 min | 72 CPU cores |
| 5. Docking of the top 100,000 REAL compounds identified in event 4 | 19 h 12 min | 72 CPU cores |
| 6. Second-round D-MPNN training | 1 h 17 min | 1 GPU, 6 CPU cores |
| 7. Generation of the second 1,000,000 compounds | 10 h 29 min | 1 GPU, 6 CPU cores |
| 8. Identification of the top 100,000 REAL compounds generated in event 7 | 6 h 36 min | 72 CPU cores |
| 9. Docking of the top 100,000 REAL compounds identified in event 8 | 18 h 54 min | 72 CPU cores |
| Total runtime (using up to 1 GPU and 72 CPU cores) | 3 d 17 h 59 min | - |

**Table S3. Numbers and proportions of REAL Space active compounds with ICM-Pro scores ≤ -30, including those sharing at least one BB with generated molecules and those with at least one better-scoring BB-sharing generated molecule, for CB2 and ROCK1.**

| Target | Actives with ICM-Pro score ≤ -30, n | Actives sharing ≥ 1 BB with generated molecules, n (%) | Actives with ≥ 1 better-scoring BB-sharing generated molecule, n (% of BB-sharing actives) |
|---|---|---|---|
| CB2 | 33 | 29 (87.9%) | 23 (79.3%) |
| ROCK1 | 48 | 38 (79.2%) | 36 (94.7%) |

**Table S4. REAL-SWIT–selected compounds evaluated for ROCK1 enzymatic activity.** Compounds with > 30% inhibition at 10 μM are shown in bold. CS-XX denotes compound XX from the previous Chemical Space Docking study[1].

| ID | Enamine ID | Chemical Structure | MW | ICM Score | Closest active compound ID | Tanimoto Similarity |
|---|---|---|---|---|---|---|
| RX-1 | Z9650641678 | 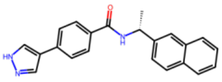 | 341.4 | -36.0 | CHEMBL3916608 | 0.73 |
| RX-2 | Z7619584337 | 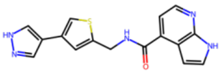 | 323.4 | -34.7 | CHEMBL1923177 | 0.48 |
| **RX-3** | Z9650641682 | 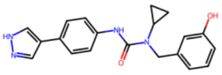 | 348.4 | -37.2 | CHEMBL2332097 | 0.63 |
| **RX-4** | Z4475094902 | 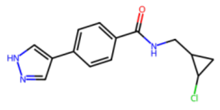 | 275.7 | -30.4 | CS-12 | 0.51 |
| RX-5 | Z3350397170 | 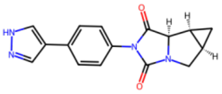 | 294.3 | -34.7 | CS-11 | 0.38 |
| **RX-6** | Z9650651395 | 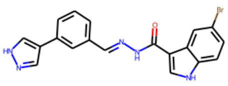 | 408.3 | -38.3 | CHEMBL1923173 | 0.52 |

| | | | | | | |
|---|---|---|---|---|---|---|
| RX-7 | Z7581771471 | 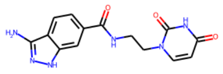 | 314.3 | -32.4 | CHEMBL1688215 | 0.36 |
| RX-8 | Z9650641696 | 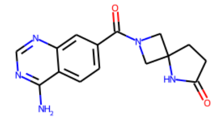 | 297.3 | -38.0 | CHEMBL1980671 | 0.28 |
| RX-9 | Z9650651499 | 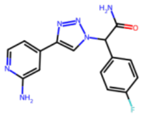 | 312.3 | -35.7 | CHEMBL3581140 | 0.35 |
| RX-10 | Z9650641700 | 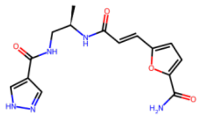 | 331.3 | -42.0 | CHEMBL3985772 | 0.31 |
| RX-11 | Z1174796555 | 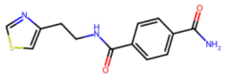 | 275.3 | -32.7 | CHEMBL3985772 | 0.33 |
| RX-12 | Z9650641713 | 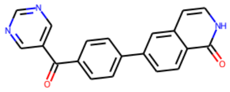 | 327.3 | -28.2 | CS-16 | 0.39 |
| **RX-13** | Z9650651938 | 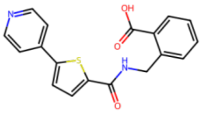 | 338.4 | -35.0 | CHEMBL1994724 | 0.52 |
| RX-14 | Z9650641720 | 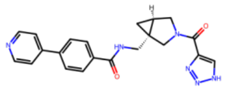 | 388.4 | -35.7 | CHEMBL1977135 | 0.44 |
| RX-15 | Z3523131090 | 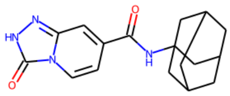 | 312.4 | -30.1 | CS-27 | 0.21 |
| RX-16 | Z9650641724 | 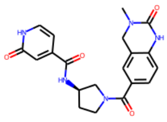 | 395.4 | -32.5 | CHEMBL3949136 | 0.34 |
| RX-17 | Z1866716423 | 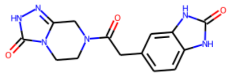 | 314.3 | -27.9 | CHEMBL3938988 | 0.28 |
| RX-18 | Z9650641727 | 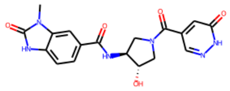 | 398.4 | -39.4 | CHEMBL3939581 | 0.30 |

| | | | | | | |
|---|---|---|---|---|---|---|
| RX-19 | Z605636572 | 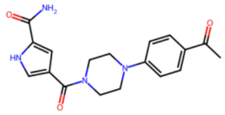 | 340.4 | -37.0 | CHEMBL1983715 | 0.34 |
| RX-20 | Z9650641728 | 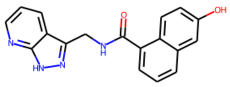 | 318.3 | -29.6 | CHEMBL4245089 | 0.43 |
| **RX-21** | Z3032829266 | 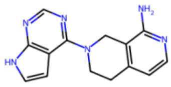 | 266.3 | -28.3 | CHEMBL591272 | 0.43 |
| RX-22 | Z7638939629 | 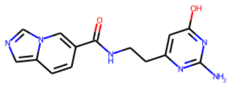 | 298.3 | -29.1 | CHEMBL3976118 | 0.32 |
| **RX-23** | Z9650641735 | 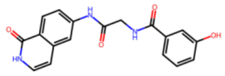 | 337.3 | -42.9 | CHEMBL1084900 | 0.46 |

### REFERENCE

1. Beroza P*, et al.* Chemical space docking enables large-scale structure-based virtual screening to discover ROCK1 kinase inhibitors. *Nature Communications* **13**, 6447 (2022).
